# MultiGEOmics: Graph-Based Integration of Multi-Omics via Biological Information Flows

**DOI:** 10.64898/2026.01.29.702593

**Authors:** Bizhan Alipour Pijani, Jubair Ibn Malik Rifat, Serdar Bozdag, the Alzheimer’s Disease Neuroimaging Initiative

**Affiliations:** Department of Computer Science & Engineering, University of North Texas, Denton, TX 76203, USA; BioDiscovery Institute, University of North Texas, Denton, TX 76203, USA; Center for Computational Life Sciences, University of North Texas, Denton, TX 76203, USA; Department of Mathematics, University of North Texas, Denton, TX 76203, USA

## Abstract

Multi-omics datasets capture complementary aspects of biological systems and are central to modern machine learning applications in biology and medicine. Existing graph-based integration methods typically construct separate graphs for each omics type and focus primarily on intra-omic relationships. As a result, they often overlook cross-omics regulatory signals—bidirectional interactions across omics layers—that are critical for modeling complex cellular processes. A second major challenge is missing or incomplete omics data; many current approaches degrade substantially in performance or exclude patients lacking one or more omics modalities. To address these limitations, we introduce **MultiGEOmics**, an intermediate-level graph integration framework that explicitly incorporates regulatory signals across omics types during graph representation learning and models biologically inspired omics-specific and cross-omics dependencies. MultiGEOmics learns robust cross-omics embeddings that remain reliable even when some modalities are partially missing. We evaluated MultiGEOmics across eleven datasets spanning cancer and Alzheimer’s disease, under zero, moderate, and high missing-rate scenarios. MultiGEOmics consistently maintains strong predictive performance across all missing-data conditions while offering interpretability by identifying the most influential omics types and features for each prediction task. The source code and the documentation of MultiGEOmics are available at https://github.com/bozdaglab/MultiGEOmics.

## 1 Introduction

Recent advances in biotechnology have made it possible to generate multi-omics data from the same samples across several diseases and conditions. Multi-omics integration strategies are commonly categorized into early, intermediate, and late integration approaches [Zitnik et al., 2019]. In early integration, features from all omics types are concatenated to form a single input representation. In intermediate integration, a joint representation is learned across omics before being passed through a shared model, as explored in [Brayet et al., 2014, Mariette and Villa-Vialaneix, 2018]. In late integration [Wang et al., 2021, Sun et al., 2018], the outputs of models trained on individual omics modalities are combined to produce the final prediction.

Several GNN-based methods proposed to integrate multi-omics data [Kesimoglu and Bozdag, 2023, Ozdemir et al., 2025]. These methods generate separate graphs for each omics modality and apply graph convolutional networks (GCNs) independently to each graph to learn omics-specific node embeddings. While this strategy effectively captures structural patterns within each omics layer, it treats each omics layer as an isolated graph during representation learning. Biological interactions across omics layers are therefore introduced only at a late integration stage, typically by fusing the learned embeddings after separate encoding. This late fusion design prevents regulatory signals from one omics layer influencing the feature learning process of another, thereby limiting the model’s ability to capture the biological interdependence and regulatory relationships that exist across omics layers.

Biological relationships within single omics layers or between individual omics layers and pathways have been studied previously [Lan et al., 2024]. In these approaches, prior biological knowledge is commonly incorporated by designing a biologically inspired hierarchical module that connects genes or microRNAs (miRNAs) to known biological pathways. However, these methods largely treat each omics layer as an independent input during representation learning. Although pathway-level features from different omics are eventually combined, the interaction between omics occurs only at a late fusion stage through simple concatenation. This modeling choice does not reflect the way biological information is organized in real systems, where molecular regulation follows a directional process in which one omics layer influences the next in a cascading manner (e.g., genomics → transcriptomics → proteomics → metabolomics) [Hasin et al., 2017]. Consequently, such approaches are limited in their ability to capture cross-omics interactions that are fundamental to biological regulation.

Although recent advances in biotechnology now allow multi-omics data to be generated, producing these data remains expensive and often constrained by experimental sensitivity. As a result, it is common for biological samples to have missing omics modalities. Such partially observed data are prevalent in real-world clinical and biomedical studies and pose a major challenge for multi-omics integration. Many existing methods are designed under the assumption that all omics modalities are available and therefore either fail when certain modalities are missing or exclude patients with incomplete data altogether, which leads to a substantial reduction in usable samples. Although several approaches have been proposed to accommodate missing omics data by learning shared latent representations or modality-specific factors [Zhao et al., 2024], their performance typically degrades substantially as the missing rate increases. This sensitivity to missing omics modalities limits the robustness and practical applicability of current multi-omics integration methods in practical settings.

To address the aforementioned limitations, we propose MultiGEOmics, a novel intermediate-level GNN-based multi-omics integration framework. MultiGEOmics explicitly incorporates biologically informed information flow into the node embedding process. Specifically, MultiGEOmics adopts a multigraph architecture, where nodes represent biological entities (e.g., patients) and edge types correspond to distinct omics modalities (e.g., genomics, epigenomics, and transcriptomics). After the first GNN message-passing layer, MultiGEOmics applies cross-omics attention among omics-specific node embeddings, with the attention structure restricted by biologically defined regulatory hierarchies and directionality. Crucially, these dependencies are not imposed as post-hoc constraints or fusion rules, but are embedded directly into the representation learning process. This design fundamentally differs from standard intermediate integration, which transforms modality-specific representations, and then the model learns a shared latent representation. In MultiGEOmics, information from upstream omics can directly influence feature learning in downstream omics during training, while reverse interactions are selectively enabled only when biologically plausible. Importantly, this cross-omics attention allows information from observed omics layers to propagate to embeddings corresponding to missing omics modalities, enabling the model to learn robust representations under complete or missing data scenarios. By incorporating attention in the representation learning process, MultiGEOmics effectively captures forward and reverse cross-omics regulatory interactions. The enriched embeddings are then propagated into the next GNN layer. Through this multigraph-based intermediate integration, MultiGEOmics achieves superior performance in downstream prediction tasks under complete data or partially missing data conditions. Extensive ablation studies further demonstrate that each model component contributes synergistically to the overall predictive performance. In summary, the key contributions of MultiGEOmics are as follows:

1. We introduce an intermediate-level integration framework that explicitly models cross-omics interactions through attention-guided information exchange across omics layers.
2. MultiGEOmics provides inherent interpretability, enabling direct biological insights, such as identifying influential omics layers and feature-level cross-omics regulatory relationships, without requiring additional explanation methods.
3. Our intermediate-level integration design allows MultiGEOmics to robustly handle missing omics data while maintaining strong predictive performance.
4. Through extensive experiments on multiple disease-related datasets, MultiGEOmics consistently outperforms existing state-of-the-art (SOTA) approaches across a wide range of evaluation metrics.

## 2 Materials and Methods

### 2.1 Preliminaries

In this section, we define the terminology and notation used to construct the graph representation from multi-omics data. The multi-omics data are represented as a multigraph 𝒢 = (𝒱, ℰ, 𝒳, 𝒜), where 𝒱 denotes the set of nodes shared across all omics modalities. The edge set is defined as ℰ = ∪_*o*∈*O*_ ℰ_*o*_, where *O* is the set of omics modalities. Each edge 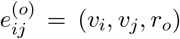 represents a connection between nodes *v*_*i*_ and *v*_*j*_ in relation type *r*_*o*_ derived from omics type *o*. The node feature matrices are denoted by 𝒳 = {*X*_*o*_ | *o* ∈ *O*}, where each 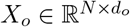 contains the node features for omics type *o*, where *N* is the number of nodes in 𝒢 and *d*_*o*_ is the corresponding feature dimension. Following common practices in multi-omics graph construction, we build an omics-specific adjacency matrix *A*_*o*_ ∈ ℝ^*N ×N*^ by computing pairwise similarities between nodes based on omics type *o* (see Supplementary Section 1 for a detailed explanation). A toy example of the resulting multi-omics graph is shown in Supplementary Figure 1. In this study, each node corresponds to a patient, and edges represent pairwise similarities between patients computed from each specific omics modality. This framework can be similarly extended to other biological entities such as genes or cell types. Under missing-data conditions, we apply the same graph-construction procedure, but treat absent omics modalities as masked inputs: when a patient lacks a given modality, all features for that modality are marked as missing.

### 2.2 Dataset

We evaluated MultiGEOmics on datasets spanning multiple diseases under both complete and missing data scenarios. The complete datasets include glioblastoma (GBM), the Religious Orders Study and Memory and Aging Project (ROSMAP), Alzheimer’s Disease Neuroimaging Initiative (ADNI), breast invasive carcinoma (BRCA), liver hepatocellular carcinoma (LIHC), bladder urothelial carcinoma (BLCA), prostate adenocarcinoma (PRAD), acute myeloid leukemia (AML), and Wilms tumor (WT). See Supplementary Section 2 for more information and the corresponding table of summary statistics. For the missing data scenario, we evaluated MultiGEOmics on the ROSMAP, BRCA, pan-kidney cohort (KIPAN), and low-grade glioma (LGG) datasets by simulating missingness. For this, we selected a proportion of patients and masked one random omics modality per each selected patient entirely, reflecting cases where a patient lacks an entire omics modality. We used masking ratios ranging from 10% to 80%. The BRCA data were obtained from two previous studies, IGCN and DeepKEGG. To distinguish between them, we refer to the dataset collected from IGCN as *TCGA_BRCA*, and the dataset collected from DeepKEGG as *BRCA*.

### 2.3 MultiGEOmics

In this section, we discuss the implementation of multi-omics information flow in MultiGEOmics and define the main architecture.

#### 2.3.1 Multi-omics information flow in MultiGEOmics

Typically, multi-omics information flow proceeds from *genomics* (e.g., SNPs, SNVs, or mutations) to *epigenomics* (e.g., DNA methylation), *transcriptomics* (e.g., mRNA expression), *proteomics*, and finally to *metabolomics*. However, certain regulatory interactions can also occur in the reverse direction between omics layers [Guo et al., 2011]. For example, miRNAs can act upstream to regulate DNA methylation [Fabbri et al., 2007]. In this study, we explicitly leverage this omics information flow to encode directional dependencies between the omics layers. Specifically, MultiGEOmics allows information to propagate across omics layers in both forward and reverse directions when learning omics-specific embeddings. For instance, in the ROSMAP dataset, which includes DNA methylation, miRNA expression, and mRNA expression modalities, the forward information flow follows *DNA methylation* → *miRNA expression* → *mRNA expression* and the reverse information flow follows *mRNA expression* → *miRNA expression* → *DNA methylation*. Supplementary Table 2 summarizes the information flow in the other datasets used in MultiGEOmics.

The next section will explain further how this information flows across omics layers and is implemented in MultiGEOmics.

#### 2.3.2 The architecture of MultiGEOmics

MultiGEOmics is a GNN-based framework in which each omics modality is handled by a separate GNN model to compute omics-specific node embeddings. Unlike existing methods, after the first GNN layer, MultiGEOmics performs two levels of attention over node embeddings: (i) intra-omics attention and (ii) cross-omics attention based on regulatory information flow between omics types. These updated embeddings are the input to the second layer of GNN. Finally, the output of the second GNN layer for each omics modality is combined and utilized for the downstream prediction task. Figure 1 provides an overview of the MultiGEOmics architecture.

**Figure 1.**
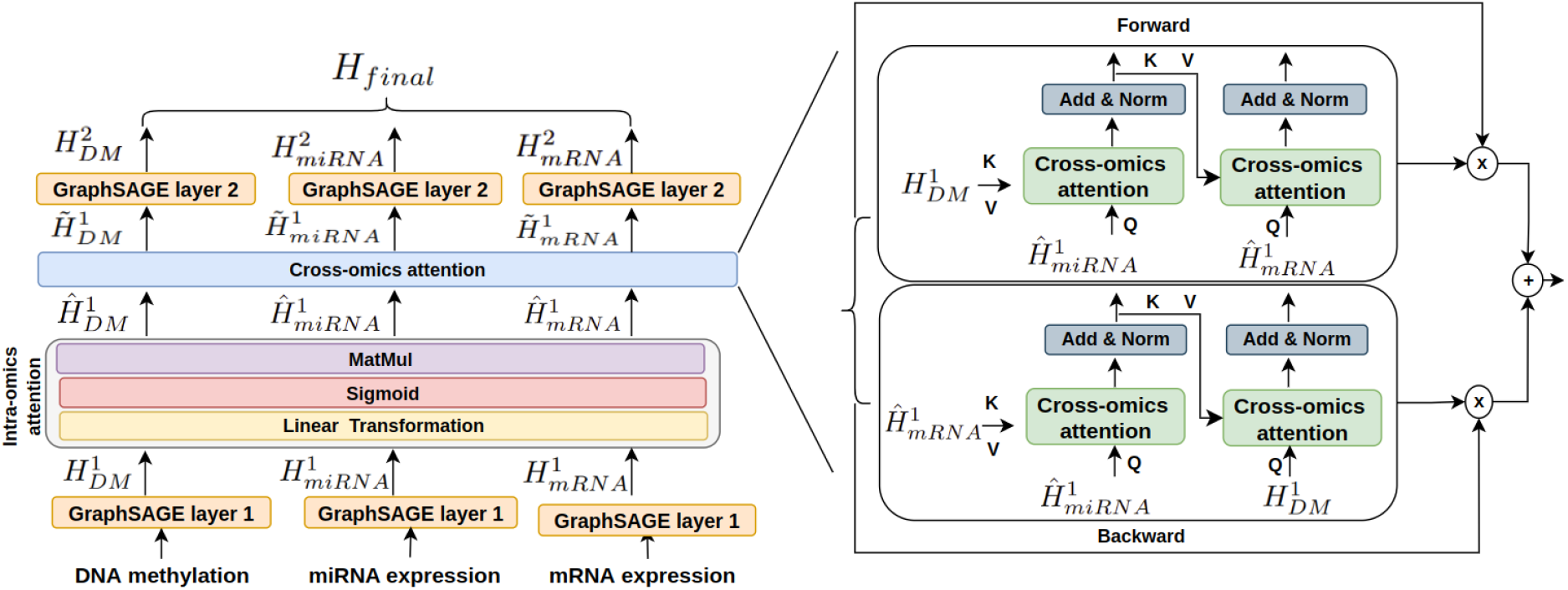
The architecture of MultiGEOmics. Each omics modality (e.g., DNA methylation, miRNA expression, and mRNA expression) is first processed by a GraphSAGE layer to learn omics-specific embeddings. Next, the cross-omics attention mechanism facilitates bidirectional information exchange between omics layers. The forward and reverse flows capture both causal and feedback regulatory relationships across modalities, while residual connections and normalization ensure stable optimization during training. The updated embeddings from this stage are passed to a second GraphSAGE layer and fused to form the final integrated representation, denoted as *H*_final_.

##### Omics-specific embeddings

We perform two hops of message passing to allow each node to aggregate information from its first- and second-hop neighbors within each omics-specific edge type, as increasing the number of hops to gather global information could lead to oversmoothing [Oono and Suzuki, 2019]. More precisely, we applied GraphSAGE [Will et al., 2017] to each omics-specific graph *G*_*o*_ to generate a representation for each node *v* specific to omics *o* as follows:

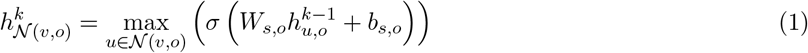

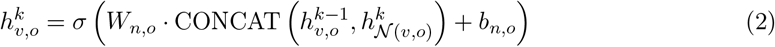

where 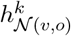 is the aggregated representation of the neighbors of node *v* in omics *o* at layer *k*, obtained using a max-pooling operation. 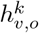 is the final embedding of node *v* at layer *k* for omics *o*, computed by combining its own previous embedding with the aggregated neighbor representation. *σ* is a non-linear activation function (e.g., ReLU) applied after the linear transformation, while CONCAT denotes the concatenation operation used to merge the transformed self-embedding 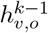 with the pooled neighbor representation 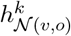. *W*_*n,o*_ and *W*_*s,o*_ are two omics-specific linear transformations, with their corresponding omics-specific bias terms *b*_*s,o*_ and *b*_*n,o*_. To update all the nodes, this process is formulated in matrix format, and we obtain *H*^*k*^ for each omics modality (e.g., 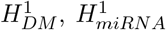, and 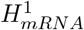) as illustrated in Figure 1 where DM stands for DNA methylation. We chose GraphSAGE as the GNN method because it enables efficient neighborhood sampling, allowing the model to scale effectively to large patient graphs. Importantly, the GNN component in MultiGEOmics is fully parameterized and modular, and it can be seamlessly replaced by alternative GNN models such as GCN and GAT [Veličković et al., 2017].

##### Intra-omics attention

For interpretability and to ensure that important features are appropriately weighted while maintaining scalability, we introduce a light-weight self-attention mechanism that produces feature-wise weights for node embeddings at the first layer for each omics modality *o* as follows:

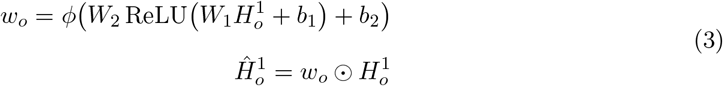

where 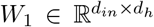 and 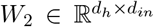 are linear transformation matrices that project the embeddings into a lower-dimensional space *d*_*h*_, and then reconstruct them back to the original dimension. *ϕ* denotes the sigmoid function used to generate feature-wise weights. These weights act as attention scores to reweight the original embeddings, resulting in omics-specific weighted embeddings 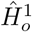 for each omics modality *o*. This process is performed independently for each omics modality.

##### Cross-omics attention

In contrast to previous works, to capture forward and reverse interactions, we allow intermediate node embeddings (e.g., 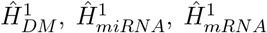) within each omics modality to interact with one another, referred to as *cross-omics attention*. We want to emphasize that cross-omics attention is not applied in an all-to-all manner. Instead, attention is defined only between biologically connected omics layers and follows a fixed directional dependency defined by biological regulation. As described in Section 2.3.1, we process the omics data using two predefined sequences: a forward biological order and its corresponding reverse order (an example from the ROSMAP dataset is shown in the Supplementary Figure 2). Under this design, attention propagates sequentially, with each omics attending to the updated representation of its predecessor. More specifically, 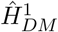 acts as key (*K*) and value (*V*), while 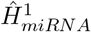 acts as query (*Q*), performing cross-omics attention between the preceding and current omics modalities, as defined below:

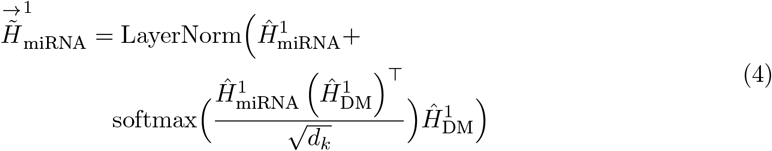

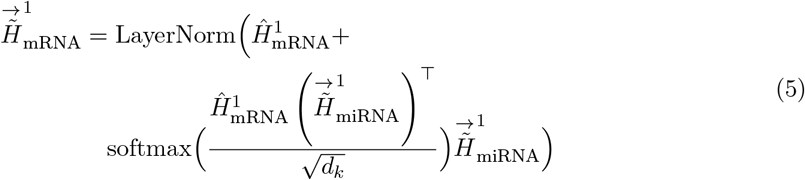

In Equations 4 and 5, miRNA and mRNA embeddings are treated as queries because these layers receive regulatory signals from their preceding omics layers. Specifically, miRNAs are influenced by DNA methylation patterns, and mRNAs are regulated by miRNAs. Therefore, these embeddings actively “ask” for relevant information from their upstream layers to update their representations. We perform the reverse pass to compute 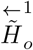 for each omics *o* (see Supplementary Section 3 for more details). We combin omics representations obtained from the forward and reverse processing orders as 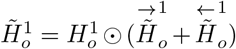 where ⊙ is Hadamard product, resulting in the updated node embeddings of the first GraphSAGE layer (e.g., 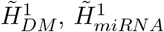 and 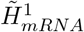). These representations are used as inputs to the second GraphSAGE layer. In this layer, only GraphSAGE message passing is applied to generate 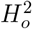 for each omics type *o* (e.g., 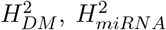, and 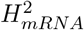) without intraomics attention or cross-omics attention. Finally, we concatenate these embeddings to form the final integrated representation, *H*_final_.

### 2.4 Learning node embeddings under missing omics scenarios

We adopt a hierarchical variational autoencoder (VAE) formulation to complete missing omics features in the learned graph embedding space. Let *M*_*o*_ ∈ {0, 1} denote the observation mask, where entries equal to 1 indicate observed features and entries equal to 0 indicate missing features for omics type *o*. The original feature matrix is masked by 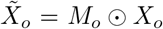. The masked features 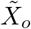 are processed using the same sequence of operations as described above. For each omics type *o*, the VAE encoder takes the output of the second GraphSAGE layer, 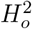, as input and maps it to a shared latent space of dimension *d* via a linear projection:

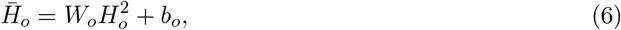

where 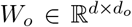 is an omics-specific trainable matrix, *d*_*o*_ is the feature dimension of 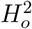, and 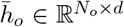. The encoder then parameterizes a Gaussian latent distribution and applies the reparameterization trick:

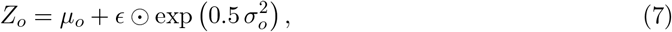

where *µ*_*o*_ and 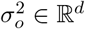 are the mean and variance for omics *o*, and *ϵ* ∼ 𝒩 (0, *I*). All omics share the same latent dimension *d*. The decoder maps the latent representation back to the original feature space of omics *o*:

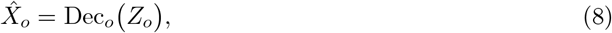

where 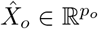 is the reconstructed omics feature vector. Because the inputs to the GraphSAGE are masked, the reconstruction loss is defined between the decoder output and the original input data, evaluated only on observed entries:

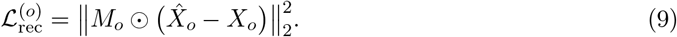

For subjects with multiple available omics, the reconstruction objective is aggregated across all omics:

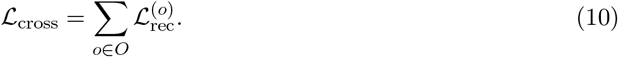

To preserve biological and structural coherence across omics modalities, we apply a hierarchically structured Kullback–Leibler (KL) divergence regularization. The first omics layer is regularized toward a standard normal prior, while the latent distribution of each subsequent layer is conditioned on the mean of the preceding modality’s latent distribution:

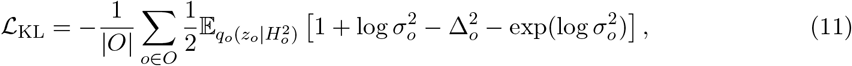

Where

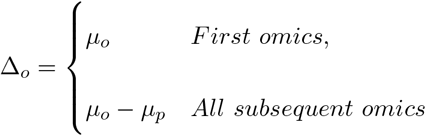

where *p* denotes the preceding omics modality with respect to *o*. For instance, *p* is miRNA when o is mRNA. Note that *order* here refers strictly to the directed information flow in the model, not to any numerical indexing of omics types. Furthermore, the hierarchical prior is forward-only: each modality *o* conditions on its immediate predecessor *p*, while no backward (reverse) conditioning is used. For the first omics modality, the KL term corresponds to the standard VAE regularization against *p*(*z*_*first*_) = 𝒩 (0, *I*), while for every subsequent modality, the conditional prior is defined as *p*(*z*_*o*_) = 𝒩 (*µ*_*p*_, *I*). This hierarchical prior structure explicitly couples latent spaces of neighboring omics modalities, enforcing progressive continuity across omics layers and preserving biologically meaningful inter-omics alignment.

### 2.5 Model loss

MultiGEOmics loss function contains different components depending on the underlying task. For datasets with missing values, we included four loss terms, whereas for complete datasets, we used two loss terms. In this study, we performed either binary or multi-class classification depending on the dataset. For example, TCGA-BRCA contains five classes, while ROSMAP has two. Accordingly, we used the cross-entropy loss function, defined as follows:

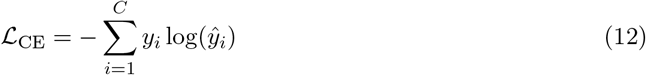

where *C* is the total number of classes, *y*_*i*_ is the true label, and *ŷ*_*i*_ is the predicted probability for class *i*. To learn meaningful patient embeddings for classification, we also applied triplet loss, which encourages the model to learn embeddings where samples from the same class are closer together, and samples from different classes are pushed further apart in the learned space. Each triplet consists of an anchor sample *s*_*a*_, a positive sample *s*_*p*_ (from the same class as the anchor), and a negative sample *s*_*n*_ (from a different class from the anchor). The triplet loss (ℒ_triplet_) is defined as follows:

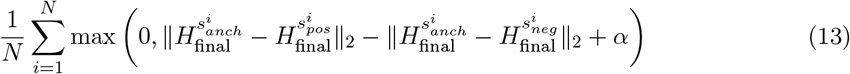

where *N* is the number of valid triplets, ∥ · ∥_2_ denotes the Euclidean (L2) norm, *α* is a margin parameter that enforces a minimum distance between the anchor-negative and anchor-positive pairs. When all omics modalities are available, the total loss is defined as ℒ = ℒ_triplet_ + ℒ_CE_. When some omics data are missing, the total loss also incorporates VAE losses, thus ℒ = ℒ_triplet_ +ℒ_CE_ +ℒ_cross_ + ℒ_KL_.

Details of the training and evaluation setup, along with hyperparameter configurations, for reproducibility, are provided in Supplementary Section 4 and Supplementary Table 3.

## 3 Results

In this section, we first present the results of MultiGEOmics across multiple multi-omics datasets. Next, we discuss the interpretation of MultiGEOmics models. Finally, we report the ablation study results to highlight the contribution of each MultiGEOmics component.

### 3.1 MultiGEOmics outperforms SOTA methods on downstream classification tasks

We benchmarked MultiGEOmics against a broad range of classification methods, including both classical models (i.e., SGD, LR, DT, KNN, SVM, RF, and MLP) and more advanced, SOTA approaches (i.e., GCN, GAT, HAN [Wang et al., 2019], HGT [Hu et al., 2020], RGCN[Schlichtkrull et al., 2018], MOGONET, SUPREME, MOGAT [Tanvir et al., 2024], HyperTMO, IGCN, Deep-KEGG, and PathCNN [Oh et al., 2021]) across eleven distinct datasets including complete and missing dataset scenarios. To ensure a fair comparison, we utilized the results reported in three prior studies (IGCN, DeepKEGG, and CLCLSA) and ran MultiGEOmics on the same train/test split.

We compared MultiGEOmics to six SOTA methods on TCGA-BRCA, TCGA-GBM, ROSMAP, and ADNI datasets reported in [Ozdemir et al., 2025]. The results show that MultiGEOmics outperformed all other SOTA models for all four datasets in all metrics except for weighted F1 in the TCGA-BRCA dataset (Table 1 and Supplementary Table 4). We also compared MultiGEOmics with eight classification methods on TCGA (BRCA, LIHC, PRAD, BLCA) and TARGET (TARGET-AML, TARGET-WT) datasets reported in [Lan et al., 2024] to predict recurrent vs. non-recurrent tumor samples. MultiGEOmics outperformed all other methods in all datasets in all evaluation metrics (Supplementary Tables 5 and 6). These results show the robustness of MultiGEOmics across multiple datasets and classification tasks.

**Table 1:** Classification results on the TCGA-BRCA, TCGA-GBM, ROSMAP, and ADNI datasets. Performance metrics for baseline methods are taken from [Ozdemir et al., 2025]. MCC: Matthews Correlation Coefficient.

| Dataset | Method | Macro F1 | MCC |
| --- | --- | --- | --- |
| TCGA-BRCA | RGCN | 0.791±0.032 | 0.744±0.027 |
|  | MOGONET | 0.765±0.027 | 0.727±0.019 |
|  | SUPREME | 0.783±0.032 | 0.742±0.031 |
|  | MOGAT | 0.806±0.016 | 0.763±0.026 |
|  | HyperTMO | 0.813±0.025 | 0.768±0.022 |
|  | IGCN | <u>0.852±0.017</u> | <u>0.821±0.014</u> |
|  | MultiGEOmics | **0.857±0.033** | **0.825±0.042** |
| TCGA-GBM | RGCN | 0.845±0.049 | 0.801±0.057 |
|  | MOGONET | 0.851±0.023 | 0.805±0.026 |
|  | SUPREME | 0.808±0.041 | 0.756±0.041 |
|  | MOGAT | 0.808±0.043 | 0.755±0.049 |
|  | HyperTMO | 0.832±0.029 | 0.781±0.034 |
|  | IGCN | <u>0.898±0.013</u> | <u>0.870±0.019</u> |
|  | MultiGEOmics | **0.944±0.032** | **0.928±0.041** |
| ROSMAP | RGCN | 0.740±0.024 | 0.503±0.049 |
|  | MOGONET | 0.781±0.019 | 0.571±0.037 |
|  | SUPREME | 0.781±0.028 | 0.575±0.056 |
|  | MOGAT | 0.757±0.031 | 0.521±0.058 |
|  | HyperTMO | 0.795±0.035 | 0.596±0.071 |
|  | IGCN | <u>0.823±0.034</u> | <u>0.659±0.069</u> |
|  | MultiGEOmics | **0.929±0.055** | **0.867±0.101** |
| ADNI | RGCN | 0.808±0.033 | 0.713±0.050 |
|  | MOGONET | 0.735±0.035 | 0.601±0.049 |
|  | SUPREME | 0.806±0.037 | 0.709±0.052 |
|  | MOGAT | 0.799±0.038 | 0.700±0.056 |
|  | HyperTMO | 0.796±0.025 | 0.695±0.042 |
|  | IGCN | <u>0.842±0.026</u> | <u>0.762±0.039</u> |
|  | MultiGEOmics | **0.973±0.016** | **0.960±0.023** |

### 3.2 MultiGEOmics maintains strong generalizability when evaluated on external held-out data

To evaluate the generalizability of MultiGEOmics across data collection sites, we conducted a centerheld-out validation on the TCGA-LIHC dataset. Specifically, patients whose sample identifiers began with the code *TCGA-DD* were excluded from the training set and used exclusively for testing. This setup ensured that all samples from one collection center (i.e., *DD*) were unseen during training, providing a scenario for external held-out evaluation. The model was trained on samples from the remaining centers using the same hyperparameter configuration as in the main experiment. Although MultiGEOmics’ performance decreased slightly when evaluated on the held-out-center data (e.g., F1 score dropped from 0.976 to 0.912) (Supplementary Table 7), it still outperformed all the SOTA methods, which were trained and tested in an easier scenario without holding out samples from a center (Supplementary Table 6). This indicates that MultiGEOmics generalizes well to samples from unseen data collection centers.

### 3.3 MultiGEOmics achieves superior performance over SOTA under missing data scenarios

To evaluate the robustness of MultiGEOmics under missing data scenarios, we followed the experimental protocol of CLCLSA [Zhao et al., 2024] and compared our model to SOTA multi-view classification methods reported in that study. Figure 2 presents the classification performance on the ROSMAP dataset across varying missing rates. As shown in the figure, MultiGEOmics consistently delivers the best performance across all three evaluation metrics for nearly all missing rates. While CLCLSA remains competitive at low missing rates, its performance drops as the amount of missing data increases. MultiGEOmics outperforms all baselines, including MVAE, SCCA, and LHGN across different missing rates, highlighting its strong robustness to missing data. The classification performance with different missing rates on LGG, KIPAN, and BRCA datasets showed the same trend (Supplementary Figure 3).

**Figure 2.**
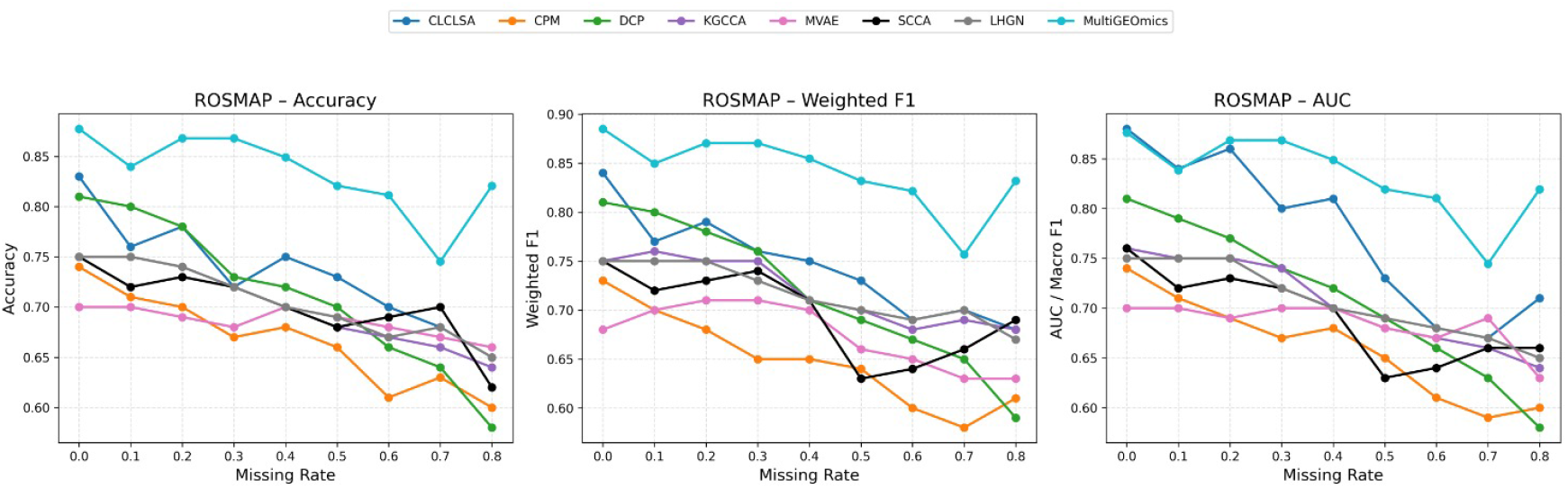
Classification performance on the ROSMAP dataset under varying feature missing rates. Results for the other methods were taken from [Zhao et al., 2024]. The ROSMAP results shown here are based on the CLCLSA benchmark dataset, which leads to differences in the results shown in Table 1 on the same dataset.

### 3.4 Interpretation of MultiGEOmics models

To interpret models trained on different datasets, we further analyzed the features identified as important by MultiGEOmics by performing literature search and functional enrichment. We also examined cross-omics attention values to find biologically relevant multi-omics associations.

#### 3.4.1 Top features of MultiGEOmics model on ROSMAP are associated with AD

To ensure robustness, we ran our model using ten different random seeds. We averaged the attention score of each feature in all ten runs and selected the top ten features with the highest attention weights from each omics type (e.g., miRNA expression, mRNA expression, and DNA methylation) for further analysis. Table 2 shows the top ten most important features for each omics type.

**Table 2:** Top ten features identified by MultiGEOmics in the ROSMAP dataset. For DNA methylation features, the corresponding genes are shown.

| DNA Methylation | miRNA expression | mRNA expression |
| --- | --- | --- |
| CD93 | ebv-miR-BART6-5p | NDE1 |
| ANGPT2;MCPH1 | hsa-miR-769-5p | ADORA2A |
| C1orf94;CSMD2 | hsa-let-7f | LINC02217 |
| CCDC8 | hsa-miR-1308 | TGFB1I1 |
| ENG | hsa-miR-100 | CMTM3 |
| RAC2 | hsa-miR-485-5p | NPNT |
| SRMS | hsa-miR-548b-3p | SLC43A3 |
| CHPT1 | mcv-miR-M1-5p | SLC4A2 |
| CDH9 | hsv1-miR-H3 | RASGRP3 |
| CCDC105 | hsa-miR-595 | ANKRD40 |

We observed that many of the identified genes have been previously linked to AD. For example, NDE1 encodes a scaffold protein essential for brain development and has been shown to regulate neurogenesis through H4K20 trimethylation–mediated heterochromatin compaction; loss of NDE1 leads to nuclear architecture defects [Chomiak et al., 2022]. Previous studies identified ADORA2A as a gene significantly associated with hippocampal volume, episodic memory performance, and cerebrospinal fluid total tau levels, highlighting its relevance to AD–related neurodegeneration [Silva et al., 2018].

Several miRNAs we identified are also known to play roles in AD. Epstein–Barr virus–derived miRNA ebv-miR-BART6-5p regulates host immune and miRNA-processing pathways through Dicer suppression. Given the growing evidence linking viral infections to AD, this miRNA may represent a potential infection-related contributor to AD pathology [Zhang et al., 2022, Hosseininasab et al., 2025]. Let-7f has been reported in prior studies to be elevated in the cerebrospinal fluid (CSF) of AD patients, suggesting a potential association with AD [Ke et al., 2025]. miR-100 is consistently differentially expressed in AD across CSF biomarker studies and tauopathy meta-analyses, and it functionally regulates amyloid-*β*–related apoptosis and microglial inflammation [Sun et al., 2021]. hsa-miR-595 is consistently upregulated in AD across serum, CSF, and brain tissues and participates in AD-related regulatory and MAPK signaling networks [Abidin et al., 2023, Öztan et al., 2025].

For the top CpG probes, we also examined the genes associated with these probes and discussed their potential biological relevance. ANGPT-2 is elevated in AD and is associated with early cerebrovascular injury, including pericyte loss and blood–brain barrier disruption. Its upregulation in both AD brains and mouse models suggests a role in early vascular dysfunction contributing to disease progression [Van Hulle et al., 2024, Flotho et al., 2025]. CSMD2 has been implicated in AD through GWAS studies in the ADNI cohort [Håvik et al., 2011, Stein et al., 2010]. Recent studies [Nasar et al., 2022, Marini et al., 2023] demonstrate that CDH9-expressing excitatory neurons in the entorhinal cortex are selectively vulnerable to AD, exhibiting early and increased tau (ptau) accumulation across Braak stages, indicating a role in early AD neurodegeneration. See Supplementary Section 6 and Supplementary Tables 9 for literature support for all selected features of ROSMAP dataset and additional gene interpretation, with the top ten features for the LIHC, PRAD, BLCA, and BRCA datasets reported in Supplementary Tables 10, 11, 12, 13, and 14, respectively.

#### 3.4.2 Functional enrichment of top MultiGEOmics features reveals disease-associated terms and pathways

We conducted enrichment analysis on the top 1% of ranked features identified by models trained on the BLCA, BRCA, LIHC, PRAD, and TCGA_BRCA datasets (Supplementary Section 7). GO enrichment in breast cancer was characterized by processes related to immune regulation and vascular-associated signaling, including negative regulation of blood pressure (GO:0045776), regulation of blood pressure (GO:0008217), and positive regulation of lymphocyte activation (GO:0051251). GO:0008217 was significantly downregulated in breast cancer, potentially reflecting dysregulation of vascular and microenvironmental processes associated with tumor progression [Xu et al., 2019]. In [Zhang et al., 2021], the authors demonstrated that BBS4 has prognostic value in patients with breast cancer. Functional enrichment analysis showed that BBS4-associated genes are involved in GO:0008217 and GO:0045776, suggesting that disruption of such regulatory signaling may be linked to breast cancer progression and patient prognosis. Additionally, GO:0051251 was significantly upregulated in the high-risk breast cancer group [Chen et al., 2023]. We showed the top GO terms and the important genes in those GO terms in the Supplementary Tables, 15, 16, 17, 18, and 19.

We performed KEGG pathway enrichment analysis to investigate the biological mechanisms associated with each dataset. Among the datasets analyzed, statistically significant KEGG pathways were identified for the TCGA_BRCA and BLCA datasets. Four KEGG pathways were significantly enriched (FDR ≤ 0.05) in TCGA_BRCA (see Supplementary Section 7 for more information). Enrichment of the PPAR signaling pathway (hsa03320) is consistent with previous findings indicating that interacting genes such as PLTP (top gene in that pathway) contribute to breast cancer tumorigenesis via PPAR pathway regulation [Chen et al., 2012]. Hormone signaling (hsa04081) is a central driver of breast cancer [Moisand et al., 2023]. Alterations in hormone signaling, together with post-transcriptional regulation by RNAs and RNA-binding proteins, contribute to breast cancer [Wang and Yang, 2021]. Enrichment of the cytokine–cytokine receptor interaction (hsa04060) pathway suggests involvement of immune and inflammatory signaling mechanisms in breast cancer [He et al., 2024].

#### 3.4.3 Attention-based discovery of cross-omics gene interactions

We investigated cross-omics attention patterns across omics layers to understand how regulatory signals propagate from genomic and epigenomic alterations to transcriptomic outcomes. Specifically, we focused on two biologically motivated omics pairs, DNA methylation/SNV–miRNA and miRNA–mRNA, with miRNAs acting as the central regulatory intermediates linking upstream genomic variation to downstream gene expression changes. We aggregated cross-omics attention values across all feature combinations and retained only the top 0.01% of interactions, thereby enriching for the most confident cross-omics dependencies. Since miRNAs act as the intermediate link between the two omics pairs, we selected only those DNA methylation/SNV–miRNA and miRNA–mRNA pairs that share the same miRNA. We particulalry focused on cases when the same gene appears simultaneously as an SNV-associated gene, a miRNA target, and an expressed mRNA, as this suggests a putative cross-omics regulatory mechanism. Using this criterion, we found 4 of 32 genes in BLCA, 5 of 34 in LIHC, 9 of 36 in PRAD, 1 of 27 genes in TCGA-BRCA, and 8 of 36 genes in BRCA. We further analyzed the these genes in each dataset to assess their relevance to the corresponding disease. Importantly, many of them have well-established roles in their respective diseases, supporting the biological relevance of the identified cross-attention interactions. See Supplementary Section 8 for more information.

### 3.5 Ablation study of key architectural components in MultiGEOmics

To assess the contribution of each architectural component in MultiGEOmics, we created several controlled variants of the model. In each variant, we modified one component at a time, for example, adding cross-omics attention after the second message-passing layer, removing cross-omics attention entirely, adding or removing reverse connections, or replacing the GraphSAGE layers with a simpler GNN such as GCN. This allowed us to isolate the effect of each architectural choice. We then evaluated all variants through an ablation study on the ADNI and TCGA_GBM datasets. In the ADNI dataset, adding a reverse connection that is not biologically supported (from lipidomics back to genomics) noticeably reduced performance (Table 3). Conversely, in the TCGA_GBM dataset, removing a biologically inspired reverse connection (from mRNA expression to miRNA expression) caused a marked drop in accuracy and F1 score, underscoring the benefit of modeling mutual omics influence when appropriate (Supplementary Table 8). Eliminating the information-sharing mechanism between omics layers consistently reduced performance, confirming that joint feature refinement enhances predictive capacity. Replacing the GraphSAGE backbone with a GCN led to performance decreases (from 0.973 to 0.850 in ADNI and 0.921 to 0.879 in TCGA_GBM), reflecting the advantage of inductive neighborhood sampling in heterogeneous biological graphs. Similarly, incorporating information sharing into the second message-passing layer slightly reduced accuracy in both datasets, suggesting that applying cross-omics attention at multiple levels can degrade performance.

**Table 3:** Ablation study on ADNI datasets.

| Method | Macro F1 | MCC |
| --- | --- | --- |
| Cross-omics attention after the second layer | 0.959 | 0.939 |
| Add reverse | 0.395 | 0.105 |
| No information sharing | 0.422 | 0.154 |
| Use GCN as backbone | 0.850 | 0.774 |
| **MultiGEOmics** | **0.973** | **0.960** |

## 4 Conclusion

We introduced MultiGEOmics, a graph-based intermediate integration framework for multi-omics learning that explicitly models cross-omics communication guided by biologically motivated information flows. MultiGEOmics learns omics-specific representations with a modular GNN backbone, then refines these embeddings using (i) intra-omics attention to emphasize informative features within each modality and (ii) cross-omics attention to propagate regulatory signals across biologically connected omics layers in forward and reverse directions. To better reflect real-world multiomics studies, MultiGEOmics is designed to operate even when some omics modalities are missing for subsets of patients.

Across eleven datasets spanning cancer and AD, MultiGEOmics outperformed SOTA methods both in no missing data and moderate-to-high missing rates scenarios. In addition, center-held-out validation demonstrated that MultiGEOmics can generalize to samples from unseen collection sites. Beyond accuracy, the attention mechanisms provide interpretability by highlighting top features and cross-omics associations.

## 6 Data availability

The data used in this study were collected from prior studies, including IGCN, DeepKEGG, and CLCLSA, and are available at https://github.com/bozdaglab/MultiGEOmics. The Alzheimer’s disease data were obtained from the ADNI database (https://adni.loni.usc.edu).

## 7 Author contributions statement

Conceptualization, B.A.P., S.B.; Methodology, B.A.P., S.B.; Data collection, B.A.P.; Running experiments, B.A.P.; Interpretation, S.B., B.A.P., J.I.M.R.; Writing-Review & Editing, S.B., B.A.P.; Visualization, B.A.P.; Supervision: S.B.

## 8 Acknowledgments

This work was supported by the National Institute of General Medical Sciences of the National Institutes of Health under Award Number R35GM133657. Data used in preparation of this article were obtained from the ADNI database (adni.loni.usc.edu). As such, the investigators within the ADNI contributed to the design and implementation of ADNI and/or provided data but did not participate in analysis or writing of this report. A complete listing of ADNI investigators can be found at: http://adni.loni.usc.edu/wp-content/uploads/how_to_apply/ADNI_Acknowledgement_List.pdf.

