## Supplemental File 1 for "MultiGEOmics: Graph-Based Integration of Multi-Omics via Biological Information Flows"

[1] Department of Computer Science and Engineering, University of North Texas
[2]BioDiscovery Institute, University of North Texas
[3]Center for Computational Life Sciences USA, University of North Texas
[4]Department of Mathematics, University of North Texas
Denton, TX 76203, USA
***Coresponding author:***
* Data used in preparation of this article were obtained from the Alzheimer's Disease Neuroimaging Initiative (ADNI) database (adni.loni.usc.edu). As such, the investigators within the ADNI contributed to the design and implementation of ADNI and/or provided data but did not participate in the analysis or writing of this report. A complete listing of ADNI investigators can be found at: http://adni.loni.usc.edu/wp-content/uploads/how_to_apply/ADNI_Acknowledgement_List.pdf.

#### 1 Graph construction

We followed the graph construction strategy introduced in the IGCN [Ozdemir et al., 2025] framework, consistent with prior work such as MOGONET [Wang et al., 2021], to construct each omics-specific graph $G_o$. Before introducing its advantages, we briefly describe the approach to ensure reproducibility and clarity.

Given an omics feature matrix $X_o \in \mathbb{R}^{N \times d_o}$, where $N$ is the number of samples (e.g., patients) and $d_o$ is the number of features for omics $o$, we first compute the cosine similarity between every pair of samples. An edge between nodes $q$ and $w$ is retained only if their similarity exceeds a threshold $\epsilon$. The adjacency matrix $A_o \in \mathbb{R}^{N \times N}$ is therefore defined as:

$$a_o^{(q,w)} = \text{Ind}(s(x_o^q, x_o^w) > \epsilon),$$

where $x_o^q$ and $x_o^w$ denote the feature vectors of nodes $q$ and $w$, respectively, and $s(x_o^q, x_o^w)$ is the cosine similarity function:

$$s(x_o^q, x_o^w) = \frac{\langle x_o^q, x_o^w \rangle}{\|x_o^q\|_2 \cdot \|x_o^w\|_2}.$$

Here, $\text{Ind}(\cdot)$ is an indicator function that returns 1 if its argument is true and 0 otherwise. The threshold $\epsilon$ is adaptively determined to maintain an average node degree $deg$ across the entire network, as defined in IGCN. Formally,

$$deg = \frac{1}{N} \sum_{q,w} \text{Ind}(s(x_o^q, x_o^w) > \epsilon).$$

We set $deg = 3$ as the default average degree.

Traditional graph construction approaches, such as $k$-nearest neighbors (kNN) or fixed similarity cutoffs, rely on manually chosen parameters (e.g., number of neighbors or distance thresholds) that strongly influence the resulting topology. These fixed-parameter methods often produce inconsistent or fragmented networks and yield variable downstream performance when applied to heterogeneous multi-omics data. In contrast, the IGCN approach adaptively determines $\epsilon$ based on the data's similarity distribution, ensuring global control over node degree and preserving only sufficiently strong biological relationships. This adaptive process leads to more robust and biologically meaningful networks, as it avoids forcing weak connections while maintaining consistent graph density across omics.

Supplementary Figure 1 illustrates a toy multi-omics graph, where the red node represents the source patient, while the black nodes are its one-hop and two-hop neighbors. The color of each edge encodes the omics type: for example, blue edges represent similarity in mRNA expression, and green edges reflect similarity based on DNA methylation.

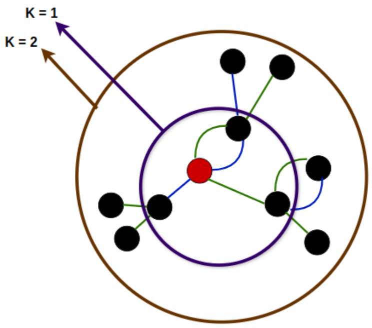


Supplementary Figure 1: An example of multi-omics graph representation with 2-hop neighbors ($k = 1, 2$).

#### 2 Datasets

We evaluated MultiGEOmics using datasets from multiple diseases, namely glioblastoma (GBM), Alzheimer's disease (AD), breast invasive carcinoma (BRCA), liver cancer (LIHC), bladder cancer (BLCA), prostate cancer (PRAD), Acute Myeloid Leukemia (AML), and Wilms Tumor (WT), Pan-kidney cohort (KIPAN), and Lower Grade Glioma (LGG). For BRCA, we used mRNA expression, DNA methylation, and miRNA expression data from TCGA to predict PAM50 subtypes, namely Basal-like, HER2-Enriched, Luminal A, Luminal B, and Normal-like. For GBM, we used mRNA expression and miRNA expression data from TCGA to predict molecular subtypes, namely Proneural, Neural, Mesenchymal, and Classical. DNA methylation was excluded due to limited sample availability.

For AD, we used mRNA expression, DNA methylation, and miRNA expression data from the ROSMAP dataset to classify AD vs. cognitively normal (CN) cases. SNPs, lipidomics, and bileomics data from the ADNI dataset were used for a three-class classification task (i.e., AD, mild cognitive impairment (MCI), and CN). We used single-nucleotide variation (SNV), mRNA expression, and miRNA expression from the TCGA Pan-Cancer Atlas to classify recurrent vs. non-recurrent tumor in BRCA, LIHC, BLCA, and PRAD. We also used mRNA expression and miRNA expression data from the TARGET database to predict recurrent vs. non-recurrent tumors in AML and WT. Supplementary Table 1 presents the statistics of all the datasets. We obtained these datasets from prior studies, including IGCN, DeepKEGG [Lan et al., 2024], and CLCLSA [Zhao et al., 2024], and used the same train/test split.

Supplementary Table 1: Description of the multi-omics datasets. CN: Cognitively Normal, MCI: Mild Cognitive Impairment, AD: Alzheimer's Disease. For the KIPAN and LGG datasets, the number of features before preprocessing is not reported; therefore, this information is listed as N/A.

| **Dataset** | **# Features before preprocessing** | **# Node features** | **# Samples (Groups)** | **# Edges** |
|---|---|---|---|---|
| **TCGA-BRCA** | mRNA expression: 20,531<br>DNA methylation: 20,106<br>miRNA expression: 503 | mRNA expression: 1,000<br>DNA methylation: 1,000<br>miRNA expression: 503 | Basal-like: 131<br>HER2-enriched: 46<br>Luminal A: 436<br>Luminal B: 147<br>Normal-like: 115 | mRNA expression: 2,629<br>DNA methylation: 2,629<br>miRNA expression: 2,631 |
| **TCGA-GBM** | mRNA expression: 12,044<br>miRNA expression: 536 | mRNA expression: 1,230<br>miRNA expression: 534 | Proneural: 164<br>Neural: 97<br>Mesenchymal: 138<br>Classical: 120 | mRNA expression: 1,559<br>miRNA expression: 1,559 |
| **ROSMAP** | mRNA expression: 55,889<br>DNA methylation: 23,788<br>miRNA expression: 309 | mRNA expression: 200<br>DNA methylation: 200<br>miRNA expression: 200 | CN: 169<br>AD: 182 | mRNA expression: 1,055<br>DNA methylation: 1,055<br>miRNA expression: 1,055 |
| **ADNI** | SNPs: 2,126,516<br>Lipidomics: 781<br>Bileomics: 24 | SNPs: 156<br>Lipidomics: 637<br>Bileomics: 24 | CN: 214<br>MCI: 210<br>AD: 178 | SNPs: 1,808<br>Lipidomics: 1,808<br>Bileomics: 1,808 |
| **PRAD** | mRNA expression: 6037<br>SNV: 4422<br>miRNA expression:409 | mRNA expression :1500<br>SNV: 2000<br>miRNA expression: 100 | Recurrent: 100<br>Non-Recurrent: 150 | mRNA expression :752<br>SNV: 870<br>miRNA expression: 752 |
| **BRCA** | mRNA expression: 6038<br>SNV: 6569<br>miRNA expression: 437 | mRNA expression:1000<br>SNV: 1000<br>miRNA expression: 100 | Recurrent: 82<br>Non-Recurrent: 129 | mRNA expression:635<br>SNV: 701<br>miRNA expression: 635 |
| **BLCA** | mRNA expression: 6025<br>SNV: 6819<br>miRNA expression: 479 | mRNA expression: 1000<br>SNV: 1000<br>miRNA expression: 100 | Recurrent: 143<br>Non-Recurrent: 259 | mRNA expression: 1208<br>SNV: 1222<br>miRNA expression:1208 |
| **LIHC** | mRNA expression: 5823<br>SNV: 5797<br>miRNA expression: 469 | mRNA expression: 1000<br>SNV: 1000<br>miRNA expression: 200 | Recurrent: 171<br>Non-Recurrent: 183 | mRNA expression: 1064<br>SNV: 1136<br>miRNA expression: 1064 |
| **TARGET-AML** | mRNA expression: 6404<br>miRNA expression: 434 | mRNA expression:2000<br>miRNA expression:100 | Recurrent: 120<br>Non-Recurrent: 101 | mRNA expression:665<br>miRNA expression:665 |
| **TARGET-WT** | mRNA expression: 6758<br>miRNA expression: 441 | mRNA expression: 2000<br>miRNA expression: 200 | Recurrent: 88<br>Non-Recurrent: 24 | mRNA expression: 338<br>miRNA expression: 338 |
| **KIPAN** | N/A | mRNA expression: 2000<br>DNA methylation: 2000<br>miRNA expression: 445 | Kidney Renal Clear Cell Carcinoma: 66<br>Kidney Renal Papillary Cell Carcinoma: 318<br>Kidney Chromophobe: 274 | mRNA expression: 1976<br>DNA methylation:1976<br>miRNA expression: 1976 |
| **LGG** | N/A | mRNA expression: 2000<br>DNA methylation: 2000<br>miRNA expression: 548 | grade-2 subjects: 246<br>grade-3 subjects: 246 | mRNA expression: 1532<br>DNA methylation:1532<br>miRNA expression: 1532 |

#### 3 Information flow

Supplementary Table 2: Forward and reverse order in the multi-omics information flow. Forward order represents the biologically guided direction of regulation (e.g., SNPs → methylation → expression). Reverse order refers to propagating information backward across omics layers. In BLCA, BRCA, LIHC, and PRAD, the lowest layer is SNV, and there is no biological justification for sending information from miRNA or mRNA expression back to SNVs; therefore, no reverse connections are used in these datasets. In the ADNI dataset—containing SNPs, bile acids, and lipids—the biological relationship between bile acids and lipids is not well established; therefore, only forward flow is used for this pair to avoid introducing noise from unsupported reverse connections.

| Dataset | Omics type | Forward | Reverse |
|---|---|---|---|
| **AML, WT, TCGA-GBM** | miRNA expression<br>mRNA expression | mRNA expression →<br>miRNA expression | miRNA expression<br>→<br>mRNA expression |
| **BLCA, BRCA, LIHC, PRAD** | SNV<br>miRNA expression<br>mRNA expression | SNV →<br>miRNA expression<br>→<br>mRNA expression | No feedback loop |
| **TCGA-BRCA** | DNA methylation<br>mRNA expression | DNA methylation →<br>mRNA expression | mRNA expression →<br>DNA methylation |
| **ADNI** | SNPs<br>bile acids<br>lipids | SNPs →<br>bile acids<br>SNPs →<br>lipids | No feedback loop |
| **KIPAN, LGG** | DNA methylation<br>miRNA expression<br>mRNA expression | DNA methylation →<br>miRNA expression<br>→<br>mRNA expression | mRNA expression<br>→ miRNA<br>expression<br>→ DNA methylation |

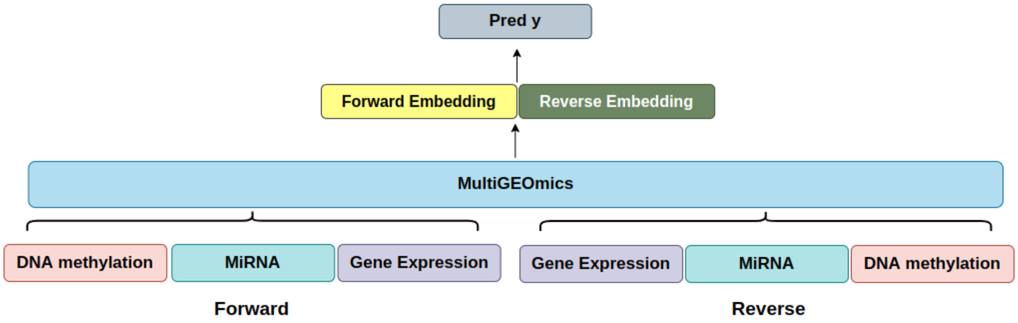


Supplementary Figure 2: Forward and reverse processing orders in MultiGEOmics. The model generates two complementary embeddings by propagating information through the omics layers in both the forward biological direction (DNA methylation → miRNA → gene expression) and the corresponding reverse direction (gene expression → miRNA → DNA methylation). These two embeddings are then fused and used for the final prediction.

To perform the reverse pass, we reverse the omics order and perform cross-omics attention as follows:

$$\overset{\leftarrow 1}{\tilde{H}}_{\text{miRNA}} = \text{LayerNorm}\Bigg(\hat{H}^1_{\text{miRNA}} + \text{softmax}\Bigg(\frac{\hat{H}^1_{\text{miRNA}}\left(\hat{H}^1_{\text{mRNA}}\right)^\top}{\sqrt{d_k}}\Bigg)\hat{H}^1_{\text{mRNA}}\Bigg) \tag{1}$$

$$\overset{\leftarrow 1}{\tilde{H}}_{\text{DM}} = \text{LayerNorm}\Bigg(\hat{H}^1_{\text{DM}} + \text{softmax}\Bigg(\frac{\hat{H}^1_{\text{DM}}\left(\overset{\leftarrow 1}{\tilde{H}}_{\text{miRNA}}\right)^\top}{\sqrt{d_k}}\Bigg)\overset{\leftarrow 1}{\tilde{H}}_{\text{miRNA}}\Bigg) \tag{2}$$

#### 4 Model training & parameters

We trained MultiGEOmics under the same condition of the other models, same data split conditions (80% training and 20% testing) with stratified sampling to preserve class balance. MultiGEOmics was optimized using the *Adamw* optimizer for 450 epochs with task-specific learning rates. To prevent overfitting, we applied early stopping during training. The process stopped if the F1 score did not improve for 150 consecutive epochs. The evaluation scores of the other methods were obtained from IGCN and DeepKEGG.

Supplementary Table 3: Hyperparameter settings for ten datasets with no missing data.

| Dataset | Optimizer | LR | Weight Decay | Hidden Emb. | Aggregator |
|---|---|---|---|---|---|
| TCGA_GBM | Adam | 0.001 | 0.0001 | 256 | pool |
| TCGA_BRCA | AdamW | 0.0001 | 0.001 | 512 | pool |
| ROSMAP | AdamW | 0.001 | 0.001 | 256 | pool |
| TARGET-WT | Adam | 0.001 | 0.001 | 256 | pool |
| TARGET-AML | Adam | 0.0002 | 0.001 | 256 | pool |
| ADNI | Adam | 0.0001 | 0.01 | 256 | pool |
| BLCA | Adam | 0.001 | 1e-3 | 512 | pool |
| LIHC | Adam | 0.0001 | 1e-3 | 256 | pool |
| PRAD | AdamW | 0.001 | 1e-3 | 256 | pool |
| BRCA | AdamW | 0.001 | 1e-3 | 256 | pool |

### 5 Supplementary Results

Supplementary Table 4: Classification results on TCGA-BRCA, TCGA-GBM, ROSMAP, and ADNI datasets. Performance metrics for baseline methods are taken from [Ozdemir et al., 2025]. MCC: Matthews Correlation Coefficient.

| Dataset | Method | Accuracy | Weighted F1 | Macro F1 | MCC |
|---|---|---|---|---|---|
| TCGA-BRCA | MLP | 0.761±0.012 | 0.752±0.015 | 0.704±0.022 | 0.648±0.020 |
| | SVM | 0.774±0.022 | 0.771±0.022 | 0.736±0.027 | 0.668±0.033 |
| | RF | 0.714±0.020 | 0.690±0.023 | 0.594±0.036 | 0.565±0.032 |
| | GCN | 0.787±0.012 | 0.782±0.016 | 0.743±0.024 | 0.685±0.020 |
| | GAT | 0.789±0.015 | 0.785±0.017 | 0.747±0.022 | 0.688±0.024 |
| | HAN | 0.781±0.025 | 0.772±0.036 | 0.714±0.069 | 0.675±0.039 |
| | HGT | 0.795±0.028 | 0.789±0.030 | 0.739±0.044 | 0.697±0.042 |
| | RGCN | 0.825±0.017 | 0.824±0.020 | 0.791±0.032 | 0.744±0.027 |
| | MOGONET | 0.813±0.013 | 0.813±0.014 | 0.765±0.027 | 0.727±0.019 |
| | SUPREME | 0.821±0.020 | 0.822±0.022 | 0.783±0.032 | 0.742±0.031 |
| | MOGAT | 0.837±0.018 | 0.839±0.017 | 0.806±0.016 | 0.763±0.026 |
| | HyperTMO | 0.838±0.015 | 0.841±0.016 | 0.813±0.025 | 0.768±0.022 |
| | IGCN | 0.874±0.014 | **0.878±0.010** | 0.852±0.017 | 0.821±0.014 |
| | MultiGEOmics | **0.875±0.029** | 0.867±0.031 | **0.857±0.033** | **0.825±0.042** |
| TCGA-GBM | MLP | 0.793±0.048 | 0.791±0.050 | 0.783±0.050 | 0.725±0.063 |
| | SVM | 0.779±0.057 | 0.777±0.056 | 0.767±0.058 | 0.706±0.075 |
| | RF | 0.818±0.049 | 0.811±0.055 | 0.801±0.059 | 0.761±0.063 |
| | GCN | 0.880±0.017 | 0.879±0.017 | 0.873±0.018 | 0.839±0.023 |
| | GAT | 0.861±0.018 | 0.859±0.019 | 0.852±0.021 | 0.814±0.025 |
| | HAN | 0.858±0.038 | 0.856±0.041 | 0.850±0.044 | 0.809±0.051 |
| | HGT | 0.840±0.033 | 0.840±0.034 | 0.839±0.036 | 0.788±0.045 |
| | RGCN | 0.851±0.043 | 0.849±0.045 | 0.845±0.049 | 0.801±0.057 |
| | MOGONET | 0.854±0.019 | 0.854±0.020 | 0.851±0.023 | 0.805±0.026 |
| | SUPREME | 0.818±0.030 | 0.816±0.034 | 0.808±0.041 | 0.756±0.041 |
| | MOGAT | 0.817±0.037 | 0.815±0.039 | 0.808±0.043 | 0.755±0.049 |
| | HyperTMO | 0.837±0.026 | 0.836±0.026 | 0.832±0.029 | 0.781±0.034 |
| | IGCN | 0.903±0.014 | 0.902±0.014 | 0.898±0.013 | 0.870±0.019 |
| | MultiGEOmics | **0.946±0.031** | **0.946±0.031** | **0.944±0.032** | **0.928±0.041** |
| ROSMAP | MLP | 0.657±0.056 | 0.650±0.059 | 0.650±0.059 | 0.335±0.111 |
| | SVM | 0.645±0.063 | 0.623±0.079 | 0.621±0.081 | 0.308±0.131 |
| | RF | 0.692±0.066 | 0.691±0.065 | 0.691±0.066 | 0.387±0.136 |
| | GCN | 0.701±0.042 | 0.700±0.042 | 0.699±0.042 | 0.405±0.085 |
| | GAT | 0.670±0.032 | 0.669±0.033 | 0.669±0.033 | 0.347±0.064 |
| | HAN | 0.775±0.025 | 0.775±0.025 | 0.774±0.025 | 0.550±0.052 |
| | HGT | 0.758±0.022 | 0.756±0.022 | 0.756±0.022 | 0.527±0.041 |
| | RGCN | 0.744±0.024 | 0.741±0.024 | 0.740±0.024 | 0.503±0.049 |
| | MOGONET | 0.782±0.019 | 0.781±0.019 | 0.781±0.019 | 0.571±0.037 |
| | SUPREME | 0.782±0.028 | 0.781±0.028 | 0.781±0.028 | 0.575±0.056 |
| | MOGAT | 0.758±0.030 | 0.757±0.031 | 0.757±0.031 | 0.521±0.058 |
| | HyperTMO | 0.796±0.035 | 0.795±0.035 | 0.795±0.035 | 0.596±0.071 |
| | IGCN | 0.824±0.034 | 0.823±0.034 | 0.823±0.034 | 0.659±0.069 |
| | MultiGEOmics | **0.930±0.055** | **0.930±0.054** | **0.929±0.055** | **0.867±0.101** |
| ADNI | MLP | 0.774±0.028 | 0.770±0.029 | 0.770±0.029 | 0.660±0.041 |
| | SVM | 0.766±0.045 | 0.763±0.048 | 0.762±0.048 | 0.651±0.065 |
| | RF | 0.791±0.047 | 0.788±0.048 | 0.788±0.049 | 0.689±0.071 |
| | GCN | 0.783±0.037 | 0.782±0.038 | 0.785±0.037 | 0.677±0.054 |
| | GAT | 0.760±0.036 | 0.759±0.035 | 0.763±0.034 | 0.642±0.055 |
| | HAN | 0.799±0.032 | 0.800±0.031 | 0.802±0.029 | 0.702±0.047 |
| | HGT | 0.758±0.039 | 0.757±0.041 | 0.762±0.041 | 0.642±0.056 |
| | RGCN | 0.807±0.034 | 0.806±0.034 | 0.808±0.033 | 0.713±0.050 |
| | MOGONET | 0.733±0.033 | 0.732±0.034 | 0.735±0.035 | 0.601±0.049 |
| | SUPREME | 0.803±0.036 | 0.803±0.038 | 0.806±0.037 | 0.709±0.052 |
| | MOGAT | 0.798±0.038 | 0.797±0.038 | 0.799±0.038 | 0.700±0.056 |
| | HyperTMO | 0.794±0.027 | 0.793±0.025 | 0.796±0.025 | 0.695±0.042 |
| | IGCN | 0.840±0.026 | 0.840±0.026 | 0.842±0.026 | 0.762±0.039 |
| | MultiGEOmics | **0.973±0.016** | **0.973±0.016** | **0.973±0.016** | **0.960±0.023** |

Supplementary Table 5: Performance comparison of methods on four TCGA datasets. Performance metrics for baseline methods are taken from [Lan et al., 2024].

| Dataset | Method | Accuracy | Recall | F1-score |
|---|---|---|---|---|
| BRCA | SGD | 0.738 ± 0.017 | 0.602 ± 0.067 | 0.631 ± 0.039 |
| | LR | 0.726 ± 0.008 | 0.501 ± 0.024 | 0.581 ± 0.015 |
| | DT | 0.580 ± 0.015 | 0.436 ± 0.014 | 0.443 ± 0.019 |
| | SVM | 0.746 ± 0.008 | 0.511 ± 0.016 | 0.604 ± 0.015 |
| | KNN | 0.615 ± 0.004 | 0.033 ± 0.014 | 0.061 ± 0.025 |
| | PathCNN | 0.488 ± 0.019 | 0.391 ± 0.038 | 0.357 ± 0.038 |
| | MOGONET | 0.763 ± 0.016 | 0.489 ± 0.034 | 0.596 ± 0.033 |
| | DeepKEGG | 0.784 ± 0.006 | 0.603 ± 0.008 | 0.672 ± 0.007 |
| | MultiGEOmics | **0.933±0.038** | **0.891±0.070** | **0.912±0.050** |
| LIHC | SGD | 0.756 ± 0.014 | 0.726 ± 0.069 | 0.740 ± 0.019 |
| | LR | 0.777 ± 0.023 | 0.732 ± 0.020 | 0.760 ± 0.023 |
| | DT | 0.522 ± 0.016 | 0.517 ± 0.026 | 0.510 ± 0.021 |
| | SVM | 0.783 ± 0.008 | 0.718 ± 0.017 | 0.762 ± 0.011 |
| | KNN | 0.555 ± 0.012 | 0.253 ± 0.014 | 0.352 ± 0.019 |
| | PathCNN | 0.521 ± 0.015 | 0.547 ± 0.033 | 0.520 ± 0.011 |
| | MOGONET | 0.812 ± 0.009 | 0.852 ± 0.026 | 0.816 ± 0.011 |
| | DeepKEGG | 0.877 ± 0.006 | 0.856 ± 0.009 | 0.870 ± 0.007 |
| | MultiGEOmics | **0.977±0.011** | **0.982±0.023** | **0.976±0.011** |
| PRAD | SGD | 0.717 ± 0.011 | 0.606 ± 0.037 | 0.622 ± 0.018 |
| | LR | 0.726 ± 0.006 | 0.530 ± 0.020 | 0.602 ± 0.017 |
| | DT | 0.613 ± 0.016 | 0.504 ± 0.018 | 0.507 ± 0.016 |
| | SVM | 0.712 ± 0.008 | 0.508 ± 0.011 | 0.581 ± 0.008 |
| | KNN | 0.613 ± 0.011 | 0.086 ± 0.025 | 0.145 ± 0.040 |
| | PathCNN | 0.556 ± 0.018 | 0.578 ± 0.077 | 0.486 ± 0.038 |
| | MOGONET | 0.647 ± 0.026 | 0.640 ± 0.030 | 0.592 ± 0.024 |
| | DeepKEGG | 0.736 ± 0.007 | 0.699 ± 0.016 | 0.679 ± 0.006 |
| | MultiGEOmics | **0.924±0.014** | **0.870±0.060** | **0.900±0.022** |
| BLCA | SGD | 0.799 ± 0.021 | 0.709 ± 0.061 | 0.712 ± 0.034 |
| | LR | 0.818 ± 0.005 | 0.668 ± 0.018 | 0.722 ± 0.011 |
| | DT | 0.567 ± 0.011 | 0.410 ± 0.012 | 0.399 ± 0.007 |
| | SVM | 0.811 ± 0.006 | 0.634 ± 0.006 | 0.703 ± 0.005 |
| | KNN | 0.621 ± 0.020 | 0.436 ± 0.080 | 0.440 ± 0.045 |
| | PathCNN | 0.567 ± 0.028 | 0.440 ± 0.039 | 0.385 ± 0.026 |
| | MOGONET | 0.777 ± 0.007 | 0.667 ± 0.033 | 0.681 ± 0.016 |
| | DeepKEGG | 0.896 ± 0.003 | 0.875 ± 0.013 | 0.857 ± 0.006 |
| | MultiGEOmics | **0.983±0.016** | **0.984±0.012** | **0.977±0.022** |

Supplementary Table 6: Performance comparison of methods on two datasets of TARGET. Performance metrics for baseline methods are taken from [Lan et al., 2024].

| Dataset | Method | Accuracy | Precision | F1-score | AUROC | AUPRC |
|---|---|---|---|---|---|---|
| TARGET-AML | SGD | 0.742 ± 0.012 | 0.785 ± 0.013 | 0.751 ± 0.014 | 0.836 ± 0.001 | 0.858 ± 0.001 |
| | LR | 0.740 ± 0.014 | 0.769 ± 0.016 | 0.758 ± 0.012 | 0.833 ± 0.012 | 0.859 ± 0.006 |
| | DT | 0.667 ± 0.015 | 0.690 ± 0.005 | 0.699 ± 0.018 | 0.661 ± 0.013 | 0.646 ± 0.009 |
| | SVM | 0.716 ± 0.014 | 0.758 ± 0.019 | 0.731 ± 0.013 | 0.784 ± 0.001 | 0.824 ± 0.004 |
| | KNN | 0.633 ± 0.008 | 0.697 ± 0.021 | 0.634 ± 0.006 | 0.667 ± 0.009 | 0.692 ± 0.007 |
| | PathCNN | 0.537 ± 0.018 | 0.571 ± 0.021 | 0.567 ± 0.017 | 0.578 ± 0.004 | 0.661 ± 0.003 |
| | MOGONET | 0.719 ± 0.014 | 0.772 ± 0.029 | 0.718 ± 0.004 | 0.806 ± 0.005 | 0.834 ± 0.007 |
| | DeepKEGG | 0.772 ± 0.005 | 0.788 ± 0.004 | 0.792 ± 0.005 | 0.850 ± 0.001 | 0.883 ± 0.001 |
| | MultiGEOmics | **0.929±0.044** | **0.940±0.056** | **0.956±0.026** | **0.968±0.021** | **0.991±0.005** |
| TARGET-WT | SGD | 0.818 ± 0.018 | 0.865 ± 0.016 | 0.887 ± 0.011 | 0.847 ± 0.009 | 0.957 ± 0.002 |
| | LR | 0.796 ± 0.016 | 0.843 ± 0.008 | 0.875 ± 0.010 | 0.844 ± 0.004 | 0.954 ± 0.005 |
| | DT | 0.673 ± 0.011 | 0.788 ± 0.008 | 0.792 ± 0.008 | 0.505 ± 0.018 | 0.789 ± 0.006 |
| | SVM | 0.787 ± 0.010 | 0.842 ± 0.007 | 0.869 ± 0.007 | 0.832 ± 0.013 | 0.953 ± 0.004 |
| | KNN | 0.791 ± 0.004 | 0.800 ± 0.003 | 0.880 ± 0.002 | 0.732 ± 0.021 | 0.882 ± 0.008 |
| | PathCNN | 0.586 ± 0.029 | 0.784 ± 0.004 | 0.708 ± 0.029 | 0.546 ± 0.027 | 0.831 ± 0.009 |
| | MOGONET | 0.823 ± 0.008 | 0.886 ± 0.018 | 0.888 ± 0.005 | 0.829 ± 0.021 | 0.948 ± 0.007 |
| | DeepKEGG | 0.830 ± 0.005 | 0.900 ± 0.006 | 0.892 ± 0.003 | 0.852 ± 0.003 | 0.962 ± 0.001 |
| | MultiGEOmics | **0.941±0.051** | **0.976±0.030** | **0.940±0.056** | **0.999±0.001** | **0.999±0.001** |

Supplementary Table 7: MultiGEOmics performance on external validation sets obtained by leaving out TCGA-LIHC collection centers during training.

| Method | Accuracy | Recall | F1-score |
|---|---|---|---|
| MultiGEOmics (with held-out-center) | 0.916±0.012 | 0.902±0.051 | 0.912±0.011 |
| MultiGEOmics (without held-out-center) | 0.977 ± 0.011 | 0.982 ± 0.023 | 0.976 ± 0.011 |

Supplementary Table 8: Ablation study on TCGA_GBM datasets.

| Dataset | Method | Accuracy | Weighted F1 | Macro F1 | MCC |
|---|---|---|---|---|---|
| TCGA_GBM | Cross-omics attention after the second layer | 0.919 | 0.919 | 0.917 | 0.892 |
| | No reverse | 0.863 | 0.864 | 0.853 | 0.816 |
| | No information sharing | 0.896 | 0.896 | 0.893 | 0.861 |
| | Use GCN as backbone | 0.879 | 0.879 | 0.874 | 0.839 |
| | **MultiGEOmics** | **0.946** | **0.946** | **0.944** | **0.928** |

Supplementary Table 9: Top 10 features identified by MultiGEOmics in the ROSMAP Dataset with supported literature. For DNA methylation features, the corresponding genes are shown.

| **DNA Methylation** | **miRNA expression** | **mRNA expression** |
|---|---|---|
| CD93 [Xu et al., 2022] | ebv-miR-BART6-5p [Zhang et al., 2022b] | NDE1 [Bradshaw et al., 2013, Chomiak et al., 2022] |
| ANGPT2;MCPH1 [Flotho et al., 2025, Chen, 2019] | hsa-miR-769-5p [Gutierrez-Tordera et al., 2024] | ADORA2A [Silva et al., 2018, Horgusluoglu-Moloch et al., 2017] |
| C1orf94;CSMD2 | hsa-let-7f [Mendes-Silva et al., 2016] | LINC02217 [Donaghy et al., 2022] |
| CCDC8 [Liu et al., 2022] | hsa-miR-1308 | TGFB1I1 [Zhang et al., 2022a] |
| ENG | hsa-miR-100 [Subasinghe et al., 2025] | CMTM3 [Vastrad and Vastrad, 2021] |
| RAC2 [Qiu and Weng, 2022] | hsa-miR-485-5p [He et al., 2021, Sun et al., 2021] | NPNT [Felsky et al., 2023] |
| SRMS | hsa-miR-548b-3p | SLC43A3 [Ou et al., 2021] |
| CHPT1 | mcv-miR-M1-5p | SLC4A2 [Kant et al., 2018] |
| CDH9 [Nasar et al., 2022, Marini et al., 2023] | hsv1-miR-H3 | RASGRP3 [Sobue et al., 2023, 2021] |
| CCDC105 [Peighan et al., 2024] | hsa-miR-595 [Öztan et al., 2025] | ANKRD40 [Li and De Muynck, 2021, Li et al., 2022] |

Supplementary Table 10: Top ten features identified by MultiGEOmics in the LIHC dataset.

| SNV | miRNA | mRNA |
|---|---|---|
| MC2R | hsa-mir-485 | TMEM97 |
| GBP2 | hsa-mir-4802 | EIF2B3 |
| SLC34A3 | hsa-mir-206 | GAS1 |
| TCF7 | hsa-mir-548k | GRPR |
| WNT10B | hsa-mir-548y | OR1F1 |
| LARS2 | hsa-mir-551b | CCNA2 |
| STAT1 | hsa-mir-210 | FBXO5 |
| MEFV | hsa-mir-767 | ZNF792 |
| REST | hsa-mir-186 | GSTM3 |
| MPO | hsa-mir-3150a | UST |

Supplementary Table 11: Top ten features identified by MultiGEOmics in the PRAD dataset.

| SNV | miRNA | mRNA |
|---|---|---|
| ABCB1 | hsa-mir-548b | EME1 |
| HYAL2 | hsa-mir-133b | DNAJC10 |
| SELL | hsa-mir-96 | SH3GLB2 |
| WT1 | hsa-mir-3173 | TYROBP |
| MPL | hsa-mir-129-1 | CCNE2 |
| CHRNA2 | hsa-mir-1269a | PPARGC1A |
| BTRC | hsa-mir-4326 | ZNF682 |
| TOB2 | hsa-mir-526b | FMO3 |
| INHBE | hsa-mir-4449 | SFTPA2 |
| ZNF124 | hsa-mir-1248 | CCNA2 |

Supplementary Table 12: Top ten features identified by MultiGEOmics in the BLCA dataset.

| SNV | miRNA | mRNA |
|---|---|---|
| CRLS1 | hsa-mir-92b | JUND |
| OCLN | hsa-mir-548l | MOCS1 |
| WDR75 | hsa-mir-4454 | PLK3 |
| UCK2 | hsa-mir-5193 | WNT3A |
| HSD17B12 | hsa-mir-4423 | DMXL1 |
| RPS6KA4 | hsa-mir-345 | SEC24C |
| FUT8 | hsa-mir-3622a | RPL3 |
| FZD5 | hsa-mir-3943 | ESPL1 |
| POFUT2 | hsa-mir-380 | LAMP2 |
| DCTN1 | hsa-mir-539 | NFKBIZ |

Supplementary Table 13: Top ten features identified by MultiGEOmics in the BRCA dataset.

| **SNV** | **miRNA** | **mRNA** |
|---|---|---|
| HMGCS2 | hsa-mir-1262 | NPR1 |
| CDC40 | hsa-mir-3188 | SLC16A10 |
| ZNF479 | hsa-mir-4745 | CKMT2 |
| GUCY2D | hsa-mir-206 | FANCG |
| PGA3 | hsa-mir-3622a | SLC36A4 |
| MGAT4B | hsa-mir-499a | SEC31B |
| CAPZB | hsa-mir-376c | PLTP |
| GDPD1 | hsa-mir-363 | FGF2 |
| DPM1 | hsa-mir-3944 | AQP8 |
| COL6A2 | hsa-mir-585 | HS6ST3 |

Supplementary Table 14: Top ten features identified by MultiGEOmics in the TCGA_BRCA Dataset. For the DNA methylation features, the corresponding genes are shown.

| **Methylation** | **miRNA** | **mRNA** |
|---|---|---|
| ARRDC2 | hsa-mir-324 | C9orf116 |
| ZFP106 | hsa-mir-92b | WWP1 |
| ASPRV1 | hsa-mir-517b | SCNN1A |
| NPSR1 | hsa-mir-3065 | GP2 |
| SLC9A9 | hsa-mir-519a-1 | POLR1E |
| C11orf92 | hsa-mir-147b | FLJ45983 |
| TDRG1 | hsa-mir-629 | TAPT1 |
| PRAME | hsa-mir-767 | UTP11L |
| SMARCB1 | hsa-mir-181c | LRRC48 |
| ADORA3 | hsa-mir-3690 | CHST2 |

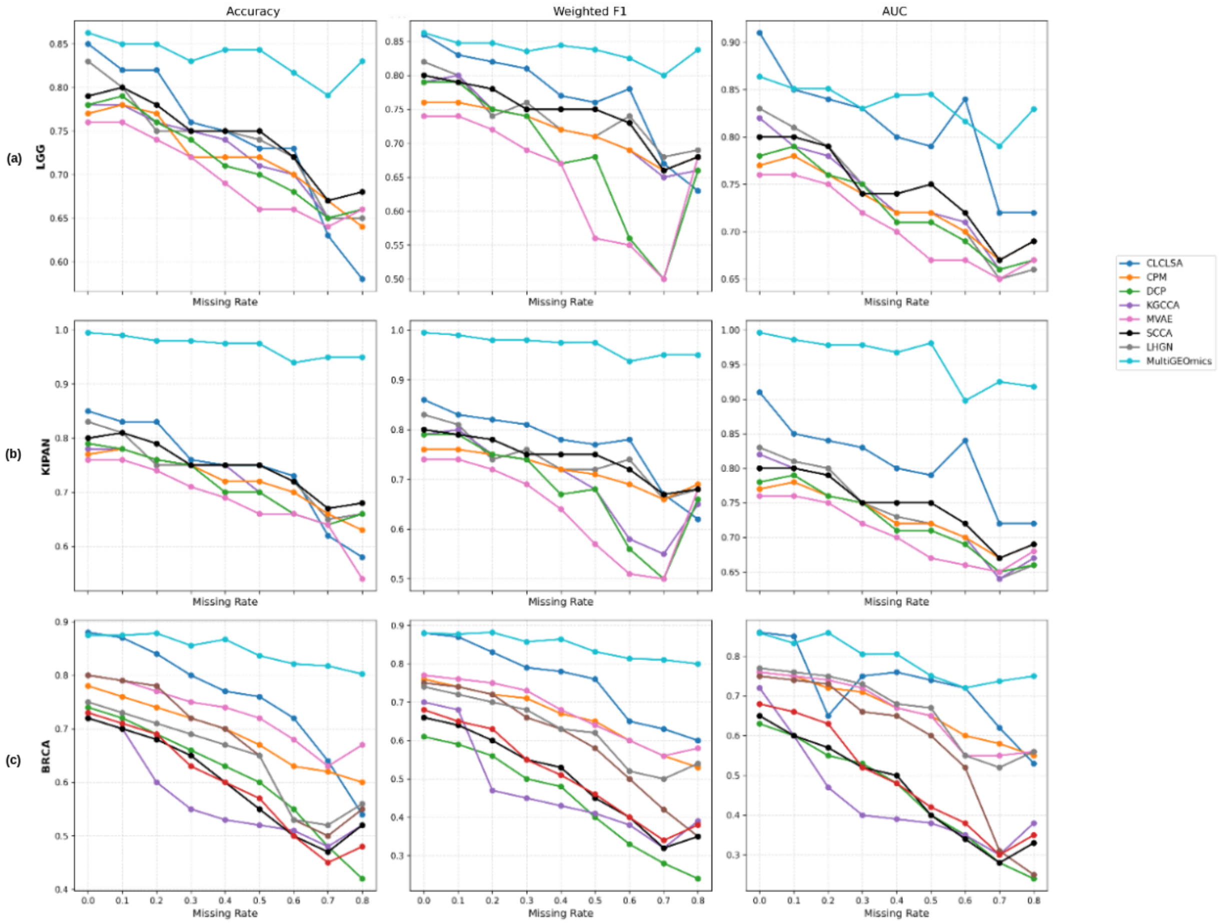


Supplementary Figure 3: Classification performance with different missing rates on a) LGG, b) BRCA, and c) KIPAN datasets. Performance metrics for baseline methods are taken from [Zhao et al., 2024].

#### 6 Literature analysis of top features in ROSMAP dataset

In [Donaghy et al., 2022], the authors combined Mild Cognitive Impairment due to AD (MCI-AD) and AD dementia into a single group for analysis. LINC02217 was significantly differentially expressed in this combined MCI-AD/AD group compared with cognitively unimpaired individuals. NPNT has been shown to play a critical role in human dental pulp stem cells and is also ranked among the top genes in large-scale AD proteomic studies [Felsky et al., 2023]. RASGRP3 was shown to be significantly downregulated in the precuneus of AD brains [Sobue et al., 2023, 2021] and was found to be upregulated in microglia in AD [Nakatsuka et al., 2025]. ANKRD40 was present in the AMP-AD upregulated gene set, showing consistent overexpression across multiple independent AD cohorts [Li and De Muynck, 2021] and was identified as a hub gene showing differential expression between normal and AD subgroups [Li et al., 2022].

#### 7 Functional Enrichment

To increase the gene set size for enrichment analysis, we constructed a unified gene list by taking the union of SNV- and mRNA-associated genes for the BLCA, LIHC, PRAD, and BRCA datasets, and the union of DNA methylation– and mRNA-associated genes for the TCGA-BRCA and ROSMAP datasets.

On the BLCA gene set, four GO terms passed the significance threshold (FDR $\leq 0.05$), including cytoplasmic translation, ribosomal small subunit biogenesis, somite development, and B cell activation (see Supplementary Table 15).

GO enrichment in BRCA revealed strong activation of neuron-related apoptotic programs, vascular and epithelial morphogenesis, hormone and steroid metabolism, immune signaling, and PI3K/MAPK pathway regulation. The most significant terms were regulation of neuron apoptotic process (GO:0043523) and neuron apoptotic process (GO:0051402) (see Supplementary Table 16).

GO enrichment analysis in LIHC revealed strong signatures of cell cycle dysregulation, DNA replication, meiosis-related programs, immune cell trafficking, and lipid metabolism—all consistent with known biological hallmarks of hepatocellular carcinoma (HCC).

On the LIHC dataset, we found the repeated enrichment of meiosis-related terms such as Meiosis I (GO:0007127), Meiotic nuclear division (GO:0140013), Meiotic cell cycle (GO:0051321, GO:1903046) (see Supplementary Table 17).

GO enrichment in PRAD was dominated by negative regulation of blood pressure(GO:0045776), positive regulation of leukocyte activation(GO:0002696), immune activation, B-cell signaling, cytokine responsiveness, blood pressure regulation, and detoxification pathways, consistent with PRAD's immunologically "cold" but cytokine-reactive microenvironment. The strongest enrichments were for positive regulation of lymphocyte activation (GO:0051251), leukocyte activation (GO:0002696), and B-cell activation (GO:0042113). Enrichment of GO:0042113 (B cell activation) is consistent with emerging evidence that activated B cells play an active role within the bladder cancer tumor microenvironment. Tumor-infiltrating $CD19^{+}$ and $CD20^{+}$ B cells have been shown to function as antigen-presenting cells, support anti-tumor T-cell responses, and are associated with improved survival and reduced recurrence in bladder cancer patients [Zhang et al., 2025].

KEGG pathway enrichment analysis of the BLCA dataset identified a highly significant enrichment of Coronavirus disease – COVID-19 (hsa05171) (see Supplementary Tables 20). Although originally annotated in the context of viral infection, this pathway captures a broad set of host cellular processes, including translation, immune signaling, and inflammatory responses, that are frequently dysregulated in cancer.

Supplementary Table 15: Significantly enriched GO biological processes for the top features in BLCA.

| GO ID | Description | Adjusted $p$-value | Top genes |
|---|---|---|---|
| GO:0002181 | Cytoplasmic translation | $6.21 \times 10^{-7}$ | RPL3, RPS11, RPL36, RPS14, RPL32, RPS7, RPL18A, RPL14, IGF2BP1, RPL13 |
| GO:0042274 | Ribosomal small subunit biogenesis | 0.0261 | RPS11, EMG1, RPS14, RPS7, GTF2H5, WDR36, PRKDC, RPS23 |
| GO:0061053 | Somite development | 0.0261 | WNT3A, IHH, PPP2R3A, LFNG, SIX1, AXIN2, PRKDC |
| GO:0042113 | B cell activation | 0.0347 | WNT3A, NFKBIZ, INHBA, FNIP1, C3, IL6, IL7, LFNG, TNFRSF13B, BLNK |

Supplementary Table 16: Significantly enriched GO biological processes for the top features in BRCA.

| GO ID | Description | Adj. $p$ | Top genes |
|---|---|---|---|
| GO:0043523 | Regulation of neuron apoptotic process | 0.003042856 | FGF2, CDC34, DKK1, CX3CL1, NTF4, CLCF1, SEMA3E, EGR1, IL6ST, EPHA4 |
| GO:0051402 | Neuron apoptotic process | 0.003042856 | FGF2, CDC34, DKK1, MAP3K5, CX3CL1, NTF4, CLCF1, SEMA3E, EGR1, IL6ST |
| GO:0035296 | Regulation of tube diameter | 0.003042856 | NPR1, ADRA2B, DRD1, CYSLTR1, HTR7, PLOD3, ADM, GPER1, SLC6A4, NTS |
| GO:0045664 | Regulation of neuron differentiation | 0.003042856 | FGF2, DKK1, ADRA2B, MEIS1, DLL1, RELN, BMP4, EPO, SHOC2, SLC6A4 |
| GO:0007188 | Adenylate cyclase-modulating GPCR signaling pathway | 0.003042856 | CACNA1D, ADRA2B, S1PR3, DRD1, HTR1F, VIPR1, HTR7, RAMP1, ADM, GPER1 |
| GO:0007204 | Positive regulation of cytosolic calcium ion concentration | 0.005064324 | S1PR3, CXCR6, CYSLTR1, ADM, CXCL13, GPER1, PTGER2, EPO, NMB, ADCYAP1R1 |
| GO:0001934 | Positive regulation of protein phosphorylation | 0.008941943 | FGF2, ADIPOQ, ADRA2B, PRLR, MAP3K5, CX3CL1, RELN, EPHA4, NBN, CTF1 |
| GO:0001666 | Response to hypoxia | 0.009654401 | ADIPOQ, ANGPTL4, IL1A, EGR1, CYSLTR1, SLC2A4, EPHA4, ADM, LIMD1, EPO |
| GO:0003018 | Vascular process in circulatory system | 0.009654401 | NPR1, ADRA2B, DRD1, CYSLTR1, HTR7, SLC2A4, SH3GL2, PLOD3, ADM, GPER1 |
| GO:0051897 | Positive regulation of PI3K/AKT signaling | 0.009654401 | FGF2, EPHA8, CX3CL1, RELN, SEMA3E, GPER1, NTS, RGL2, NTRK3, NTRK2 |

Supplementary Table 17: Significantly enriched GO biological processes for the top features in LIHC.

| **GO ID** | **Adjusted $p$-value** | **Description** | **Top genes** |
|---|---|---|---|
| GO:0007127 | 0.0126 | Meiosis I | FBXO5, BCL2L11, CCNE1, NDC1, CKS2, CDC25B, BRIP1, STAG3, CCNE2 |
| GO:0006261 | 0.0126 | DNA-templated DNA replication | FBXO5, EXO1, CCNE1, ORC1, BARD1, CDC7, POLD3, RFC5, POLE3, CCNE2 |
| GO:0051321 | 0.0126 | Meiotic cell cycle | FBXO5, BCL2L11, EXO1, CCNE1, LFNG, NDC1, CKS2, CDC25B, BRIP1, SMC1A |
| GO:0061982 | 0.0126 | Meiosis I cell cycle process | FBXO5, BCL2L11, CCNE1, NDC1, CKS2, CDC25B, BRIP1, STAG3, CCNE2 |
| GO:0140013 | 0.0301 | Meiotic nuclear division | FBXO5, BCL2L11, CCNE1, NDC1, CKS2, CDC25B, BRIP1, STAG3, CCNE2, PLCB1 |
| GO:0002691 | 0.0312 | Regulation of cellular extravasation | IL27RA, SELP, SELE, CCL21, PLCB1 |
| GO:0051168 | 0.0312 | Nuclear export | STRADB, BARD1, NUP155, THOC7, RAN, PHAX, CDKN2A, EIF4E, NUP62 |
| GO:0046513 | 0.0312 | Ceramide biosynthetic process | UGCG, DEGS2, GAL3ST1, DEGS1, CERS5, B4GALNT1 |
| GO:0071900 | 0.0312 | Regulation of protein serine/threonine kinase activity | CDK5R1, FGF1, IPO7, MST1R, IRAK3, CD24, NTF3, CDKN2A, PAK1, NUP62 |
| GO:1903046 | 0.0312 | Meiotic cell cycle process | FBXO5, BCL2L11, CCNE1, NDC1, CKS2, CDC25B, BRIP1, STAG3, CCNE2, PLCB1 |

Supplementary Table 18: Significantly enriched GO biological processes for the top features in RPAD.

| GO ID | Adjusted $p$-value | Description | Top genes |
|---|---|---|---|
| GO:0045776 | 0.0023 | Negative regulation of blood pressure | SOD2, TAC1, ABAT, UCN, NOS2, NOS3, BDKRB1, GLP1R |
| GO:0051251 | 0.0023 | Positive regulation of lymphocyte activation | TYROBP, SIRPB1, FCGR3A, SYK, CLCF1, LILRB4, MAD2L2, TNFSF13B, IL18, STAT6 |
| GO:0008217 | 0.0023 | Regulation of blood pressure | PPARG, SOD2, SERPINF2, ADH5, TAC1, COL1A2, ABAT, UCN, CYP4F2, UTS2B |
| GO:0002696 | 0.0060 | Positive regulation of leukocyte activation | TYROBP, SIRPB1, FCGR3A, SYK, CLCF1, LILRB4, MAD2L2, TNFSF13B, IL18, STAT6 |
| GO:0050864 | 0.0060 | Regulation of B cell activation | TYROBP, CASP3, SYK, CLCF1, MAD2L2, TNFSF13B, STAT6, EPHB2, INHBA, TCF3 |
| GO:0043409 | 0.0094 | Negative regulation of MAPK cascade | GPS2, PPARG, PRMT1, SMAD4, EPHA4, HYAL2, LILRB4, SH2B3, EPHB2, NLRP12 |
| GO:0070663 | 0.0115 | Regulation of leukocyte proliferation | TYROBP, ENPP3, FCGR3A, TNFRSF1B, CASP3, SYK, CLCF1, LILRB4, CSF1R, TNFSF13B |
| GO:0002237 | 0.0115 | Response to molecule of bacterial origin | ZFP36, SOD2, ADH5, TNFRSF1B, E2F1, CASP3, HMGCS2, CD68, NR4A1, CD14 |
| GO:0042113 | 0.0156 | B cell activation | TYROBP, GPS2, EZH2, CASP3, LYL1, SYK, CLCF1, MAD2L2, TNFSF13B, STAT6 |
| GO:0032496 | 0.0156 | Response to lipopolysaccharide | ZFP36, SOD2, ADH5, TNFRSF1B, E2F1, CASP3, HMGCS2, CD68, NR4A1, CD14 |

Supplementary Table 19: Significantly enriched GO biological processes for the top features in TCGA_BRCA.

| **GO ID** | **Description** | **adj. *p*-value** | **Top genes** |
|---|---|---|---|
| GO:0045776 | Negative regulation of blood pressure | 0.0023 | SOD2, TAC1, ABAT, UCN, NOS2, NOS3, BDKRB1, GLP1R |
| GO:0051251 | Positive regulation of lymphocyte activation | 0.0023 | TYROBP, SIRPB1, FCGR3A, SYK, CLCF1, LILRB4, MAD2L2, TNFSF13B, IL18, STAT6 |
| GO:0008217 | Regulation of blood pressure | 0.0023 | PPARG, SOD2, SERPINF2, ADH5, TAC1, COL1A2, ABAT, UCN, CYP4F2, UTS2B |
| GO:0002696 | Positive regulation of leukocyte activation | 0.0060 | TYROBP, SIRPB1, FCGR3A, SYK, CLCF1, LILRB4, MAD2L2, TNFSF13B, IL18, STAT6 |
| GO:0050864 | Regulation of B cell activation | 0.0060 | TYROBP, CASP3, SYK, CLCF1, MAD2L2, TNFSF13B, STAT6, EPHB2, INHBA, TCF3 |
| GO:0050867 | Positive regulation of cell activation | 0.0086 | TYROBP, SIRPB1, FCGR3A, SYK, CLCF1, LILRB4, MAD2L2, TNFSF13B, IL18, STAT6 |
| GO:0043409 | Negative regulation of MAPK cascade | 0.0094 | GPS2, PPARG, PRMT1, SMAD4, EPHA4, HYAL2, LILRB4, SH2B3, EPHB2, NLRP12 |
| GO:0070663 | Regulation of leukocyte proliferation | 0.0115 | TYROBP, ENPP3, FCGR3A, TN-FRSF1B, CASP3, SYK, CLCF1, LILRB4, CSF1R, TNFSF13B |
| GO:0002237 | Response to molecule of bacterial origin | 0.0115 | ZFP36, SOD2, ADH5, TNFRSF1B, E2F1, CASP3, HMGCS2, CD68, NR4A1, CD14 |
| GO:0042113 | B cell activation | 0.0156 | TYROBP, GPS2, EZH2, CASP3, LYL1, SYK, CLCF1, MAD2L2, TNFSF13B, STAT6 |
| GO:0050871 | Positive regulation of B cell activation | 0.0191 | SYK, CLCF1, MAD2L2, TNFSF13B, STAT6, EPHB2, TCF3, SPI1 |

Supplementary Table 20: Significantly enriched KEGG pathways for top genes in BLCA.

| **KEGG ID** | **Description** | **adj. *p*-value** | **Top genes** |
|---|---|---|---|
| hsa05171 | Coronavirus disease - COVID-19 | 0.0020 | RPL3, RPS11, C3, IL6, RPL36, RPS14, RPL32 |

Supplementary Table 21: Significantly enriched KEGG pathways for top genes in BRCA.

| Pathway ID | Pathway Name | Adjusted P-value | Top Genes |
|---|---|---|---|
| hsa03320 | PPAR signaling pathway | 0.01679 | PLTP, ADIPOQ, PLIN1, ANGPTL4, ILK, ME1, PLIN4 |
| hsa04081 | Hormone signaling | 0.01679 | NPR1, SLC16A10, ADIPOQ, ADRA2B, PRLR, HEPHL1, DRD1 |
| hsa04080 | Neuroactive ligand-receptor interaction | 0.01679 | ADRA2B, GRPR, S1PR3, RLN1, PRLR, CHRNA1, GLRA3 |
| hsa04060 | Cytokine-cytokine receptor interaction | 0.02503 | IL1RL2, PRLR, CX3CL1, CLCF1, IL1A, IFNL2, CXCR6 |

#### 8 Cross-omics attention visualization

In the BLCA dataset, genes such as COL11A1 [Li et al., 2025], ALDH1L1 [Meyers et al., 2025], and ACADVL [Xiong et al., 2022] have been reported to be associated with bladder cancer (Supplementary Figure 4). This suggests that coordinated genomic, epigenomic, and transcriptomic regulation contributes to bladder cancer pathology.

In the LIHC dataset, VPS35 [Zhang et al., 2020] and SOX4 [Wang et al., 2017], both known to influence oncogenic signaling and liver cancer progression, are identified as shared targets across omics layers (Supplementary Figure 5).

In the PRAD dataset, multiple genes highlighted by cross-attention analysis, including IL6ST [Sternberg et al., 2024], FMOD [Silva et al., 2022], TAOK3 [Romanuik, 2008], SH2B2 [Marzec et al., 2021], KLC2 [Wang et al., 2015], RASAL3 [Mishra et al., 2019] have been previously associated with prostate cancer (Supplementary Figure 6). The emergence of these genes suggests that prostate cancer progression may involve multi-layer regulatory coordination.

Likewise, in the BRCA datasets, genes such as SGPP1 [Nema and Kumar, 2021], TAPT1 [Dang et al., 2020], ABCA13 [Krøigård et al., 2018], CHST2 [Liu et al., 2025], GPR161 [Feigin et al., 2014], and BTG3 [Yu et al., 2018] are highlighted as cross-omics convergence points (Supplementary Figures 7 and 8). These findings indicate that breast cancer phenotypes may arise from cascading regulatory effects across omics layers, rather than isolated alterations at a single molecular level.

These results demonstrate that the learned cross-omics capture biologically meaningful, disease-relevant regulatory programs, supporting the interpretability and translational relevance of our model.

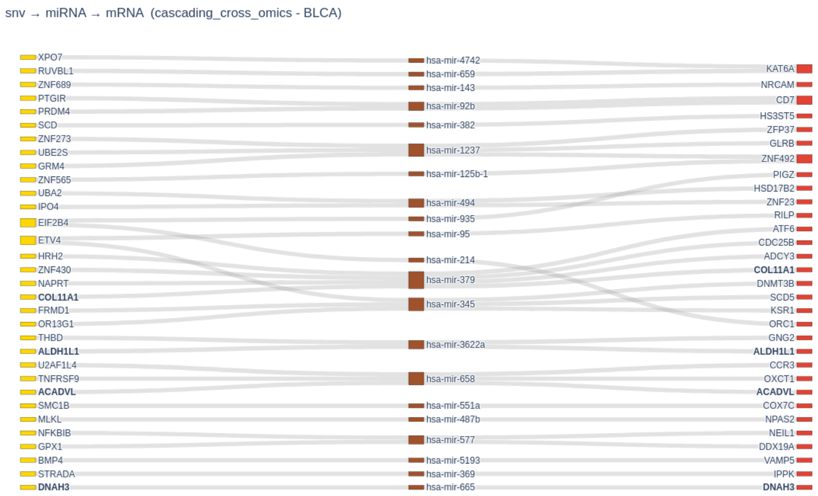


Supplementary Figure 4: BLCA cross-omics attention Sankey plot. Genes that appear in SNV and mRNA layers are shown in bold, as these genes might be involved in a cross-omics regulatory pathway.

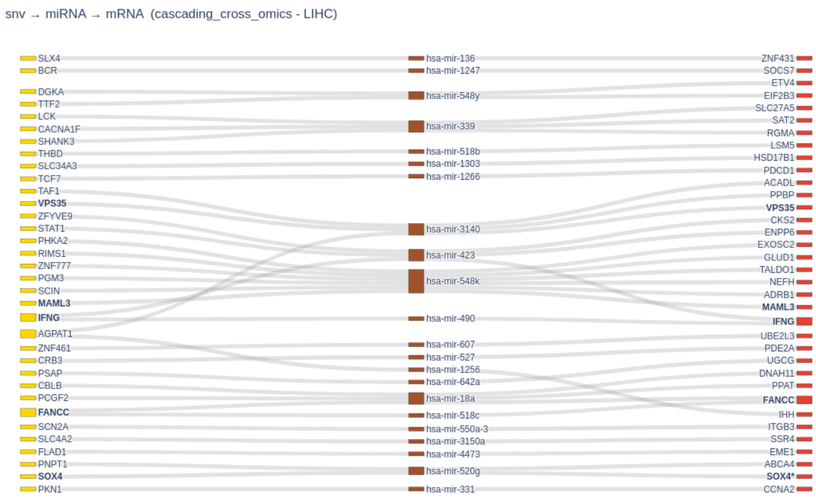


Supplementary Figure 5: LIHC cross-omics attention Sankey plot. Genes that appear in SNV and mRNA layers are shown in bold, as these genes might be involved in a cross-omics regulatory pathway.

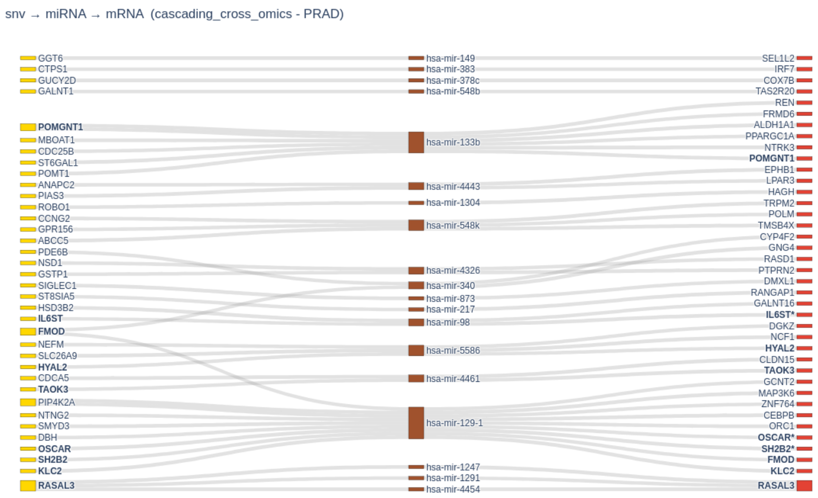


Supplementary Figure 6: PRAD cross-omics attention Sankey plot. Genes that appear in SNV and mRNA layers are shown in bold, as these genes might be involved in a cross-omics regulatory pathway.

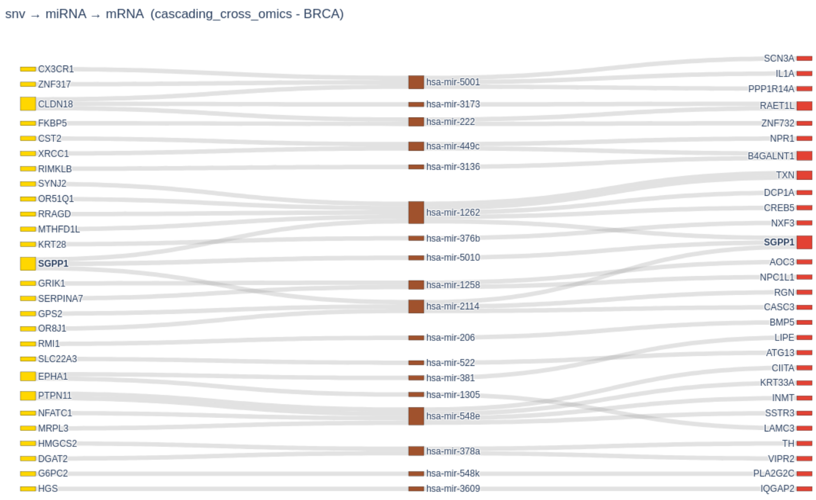


Supplementary Figure 7: BRCA cross-omics attention Sankey plot. Genes that appear in SNV and mRNA layers are shown in bold, as these genes might be involved in a cross-omics regulatory pathway.

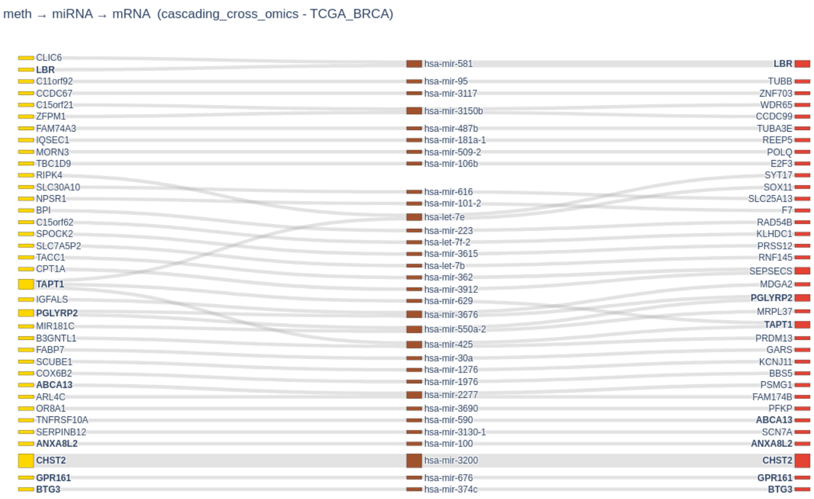


Supplementary Figure 8: TCGA_BRCA cross-omics attention Sankey plot. Genes that appear in SNV and mRNA layers are shown in bold, as these genes might be involved in a cross-omics regulatory pathway.
